# The translatome of adult cortical axons is regulated by learning *in vivo*

**DOI:** 10.1101/502419

**Authors:** Linnaea E. Ostroff, Emanuela Santini, Robert Sears, Zachary Deane, Joseph E. LeDoux, Tenzin Lhakhang, Aristotelis Tsirigos, Adriana Heguy, Eric Klann

## Abstract

Local translation can support memory consolidation by supplying new proteins to synapses undergoing plasticity. Translation in adult forebrain dendrites is an established mechanism of synaptic plasticity and is regulated by learning, yet there is no evidence for learning-regulated protein synthesis in adult forebrain axons, which have traditionally been believed to be incapable of translation. Here we show that axons in the adult rat amygdala contain translation machinery, and use translating ribosome affinity purification (TRAP) with RNASeq to identify mRNAs in cortical axons projecting to the amygdala, over 1200 of which were regulated during consolidation of associative memory. Mitochondrial and translation-related genes were upregulated, whereas synaptic, cytoskeletal, and myelin-related genes were downregulated; the opposite effects were observed in the cortex. Our results demonstrate that learning-regulated axonal translation occurs in the adult forebrain, and support the likelihood that local translation is more a rule than an exception in neuronal processes.

## Introduction

Neurons use local translation as a means of rapid, spatially-restricted protein regulation in their distal processes, particularly during remodeling driven by external cues ^1–3^. Memory consolidation requires new proteins to stabilize molecular changes induced by learning^4,5^, and local translation in dendrites is thought to be an essential source of these proteins^6^. Rich and diverse assortments of mRNAs have been described in neuropil of the mature hippocampus^7–9^ and in cortical synaptoneurosomes^10^, underscoring the importance of decentralized translation in synaptic function. Yet no role for axonal translation in learning and memory has been reported in the adult forebrain.

Translation has long been known to occur in invertebrate axons, and it is now established to be essential for growth and response to guidance cues in developing CNS axons, and in regeneration of PNS axons ^11–14^. Adult forebrain axons, in contrast, traditionally have been characterized as lacking the capacity for translation, in part due to a lack of reliable evidence, and in part to the perception that they are structurally and functionally inert compared to dendrites and immature axons.^11,12,15^. However, a number of recent studies have shown that mature axons are in fact capable of translation, at least in some circumstances ^16–18^, including in the CNS ^19–22^. This work has largely been done with cultured neurons, but one study used translating ribosome affinity purification (TRAP) to isolate ribosome-bound mRNAs in retinal ganglion cells (RGCs) of adult mice ^20^, demonstrating that translation does occur in adult CNS axons *in vivo*. Presynaptic translation has been shown to be necessary for long-term depression in hippocampal^23^ and striatal^24^ slice preparations from young animals, indicating that axonal translation is involved in synaptic plasticity and therefore could be important in memory as well.

Auditory Pavlovian conditioning (fear or threat conditioning), in which animals learn to associate an auditory tone with a foot shock, is supported by persistent strengthening of synaptic inputs to the lateral amygdala (LA) from auditory areas ^25^. The LA receives strong excitatory input from auditory cortical area TE3^26–28^, and Pavlovian conditioning induces persistent enhancement of presynaptic function at these synapses^29,30^. Consolidation of threat memory requires translation in the LA ^31^, and we have found that it induces changes in the translational machinery in LA dendrites associated with synapse enlargement^32^. Intriguingly, we also found that learning-induced structural changes occurred at individual axonal boutons as opposed to uniformly along axons, suggesting that plasticity may be as synapse specific and compartmentalized on the presynaptic side as it is on the postsynaptic side^33^. To determine whether axonal translation is involved in memory formation, we confirmed the presence of translation machinery in LA axons, and combined TRAP with RNAseq to identify changes in the translatome of auditory cortical axons during memory consolidation.

### Adult axons contain translation machinery

Early electron microscopy studies reported abundant polyribosomes in the somata and dendrites of neurons, but rarely in axons (reviewed in ^12,34^). However, the puacity of conspicuous polyribosomes does not necessarily preclude translation. Regenerating sciatic nerve axons contain mRNAs and translate membrane proteins *in vivo*, but do not show ultrastructural evidence of polyribosomes or rough endoplasmic reticulum (ER)^35,36^. In addition, hippocampal interneuron axons contain ribosomal proteins ^23^. This suggests that translation sites other than the classic morphological structures do exist, such as the periaxoplasmic ribosomal plaques found in adult spinal cord axons^37^. Recent work in yeast has shown that translation can occur on 80S monosomes, with a bias towards highly regulated transcripts ^38^.

We have occasionally observed polyribosomes in presynaptic boutons in the adult rat LA by EM (Figure 1a-b, Supplementary Figure 1a-e), although these are infrequent (LO, unpublished observations). A possible explanation for this is that these axons contain translation machinery that does not usually assemble into polyribosomal structures with traditionally recognizable morphology. To more directly assess the potential for translation in LA axons, we used immuno-electron microscopy to localize components of the translation machinery. Because translation initiation is most extensively regulated step in gene expression, as well as a critical mediator of memory formation^39^, we focused on translation initiation factors. The eukaryotic initiation factors eIF4E, eIF4G, and eIF2α each were present in axons forming synapses onto spiny dendrites in the caudal dorsolateral subdivision of the LA (Figure 1c-e), which receives the most robust projections from TE3^26–28^, as was ribosomal protein S6 (Figure 1f). These synapses have the same classic excitatory morphology as the glutamatergic projections from TE3 to LA^28^, consistent with local translation on TE3 inputs. Quantification of eIF4E immunolabel through serial sections of neuropil revealed that 63% of axons were labeled, along with 39% of dendritic spines and 100% of dendritic shafts (Supplementary Figure 1f-i). Consistent with this pattern, we have previously found polyribosomes throughout dendritic shafts but in only a subset of dendritic spines, where their presence is regulated by learning^32^.

**Figure 1.**
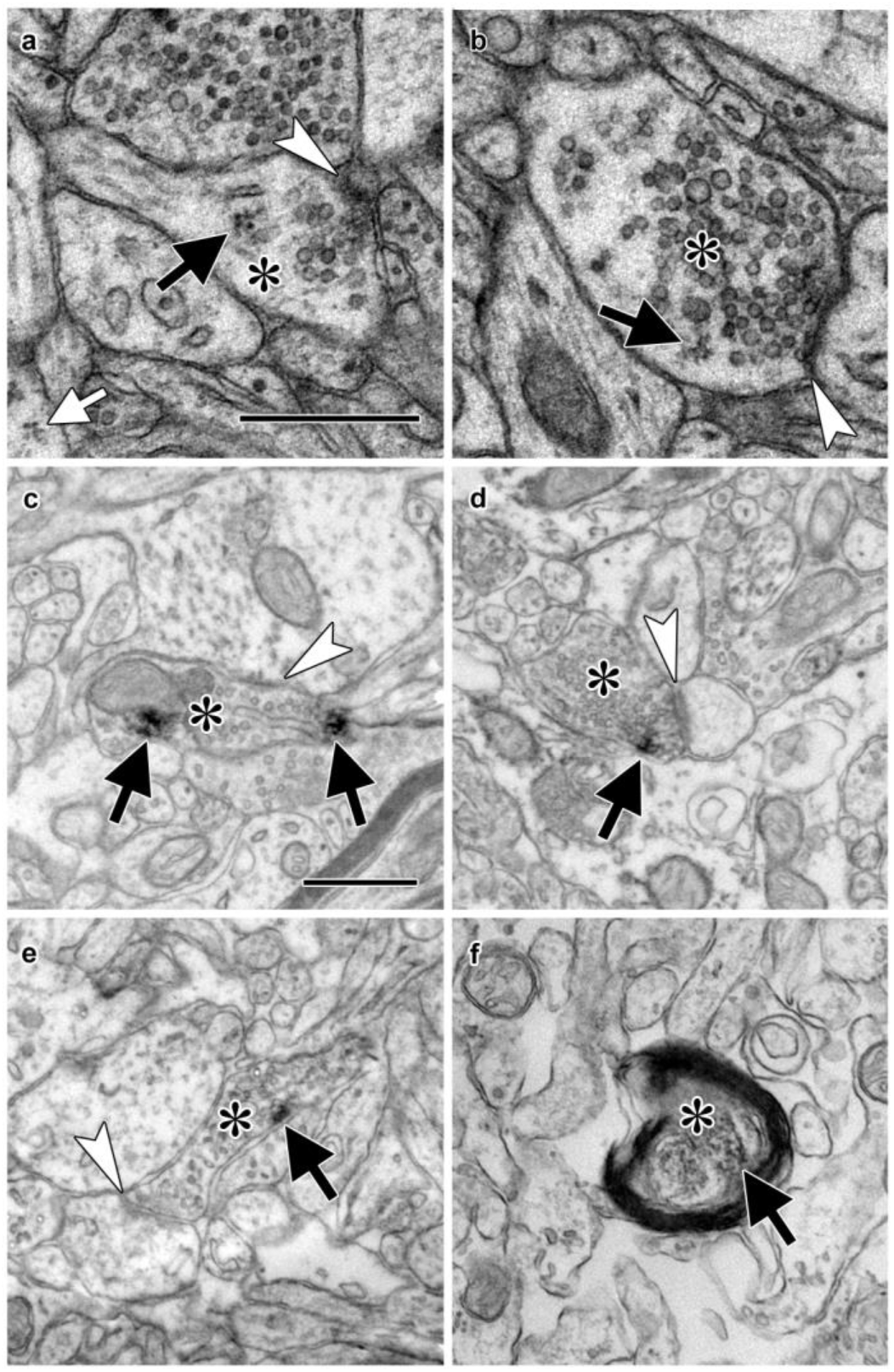
Electron micrographs of translation machinery in lateral amygdala axons. a-b) Polyribosomes (black arrows) in axonal boutons (asterisks). A polyribosome in an astrocytic process (white arrow) is visible at the lower left of panel (a). c-e) Axonal boutons (asterisks) containing immunolabeling (black arrows) for eIF4E (c), eIF4G1 (d), and eIF2α (e). White arrowheads indicate asymmetric synapses onto dendritic spines (a, d, and e) and shafts (b and c). f) Myelinated axon (asterisk) containing immunolabel for ribosomal protein s6 (arrow). Scale bars = 500nm.

### Isolation of the adult axonal translatome

To identify mRNA transcripts translated in distal TE3 axons, we used TRAP^40^, in which a tagged ribosomal protein is expressed in cells of interest and used to immunoprecipitate ribosome-bound mRNA. A recent study used an HA-tagged ribosomal protein to examine the translatome of retinal ganglion cell axons in both immature and adult mice^20^, and an eGFP-tagged ribosomal protein expressed in adult mouse layer V cortical neurons was observed in axons of the corticospinal tract^41^, demonstrating that this method is viable in at least two types of adult CNS neurons *in vivo*. We used a viral vector to express an eYFP-ribosomal protein L10a fusion protein^42^ in TE3 cells in adult rats (Figure 2a-b). Pilot experiments using an adeno-associated viral vector resulted in moderate to strong retrograde infection of cells in afferent areas. To ensure that no cell bodies outside of the injection site expressed the construct, we switched to a lentiviral vector, which did not result in retrograde infection. Immuno-electron microscopy confirmed the presence of eYFP in LA axons (Figure 2c-f).

**Figure 2.**
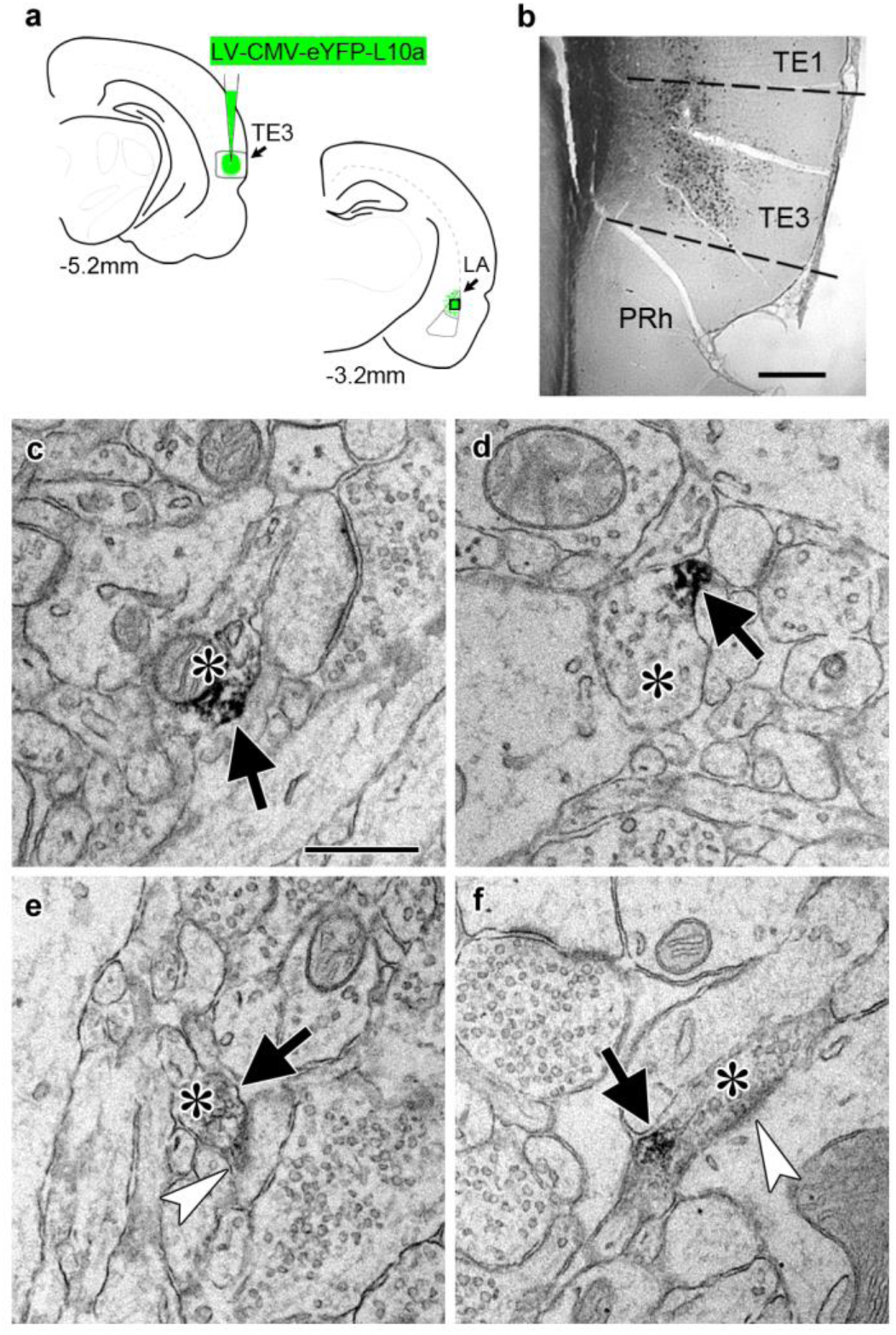
Transport of a tagged ribosomal L10a protein to cortical projection axons. a) Schematic of injection site in cortical area TE3 and its lateral amygdala (LA) projection area, with AP coordinates from Bregma noted. The black square indicates the area of LA sampled for EM. PRh: perirhinal cortex. b) Immunolabeling of YFP in transfected TE3. c-f) Electron micrographs of LA showing axonal boutons (asterisks) containing YFP immunolabel (black arrows). The boutons in (e) and (f) are forming asymmetric synapses (white arrowheads) on a dendritic spine head (e) and a dendritic shaft (f). Scale bars=500µm in (b) and 500nm in (c-f).

TRAP was combined with Pavlovian conditioning to determine how the axonal translatome changes during memory consolidation (Figure 3a). Animals expressing eYFP-L10a in TE3 were given either Pavlovian conditioning, consisting of auditory tones paired with mild foot shocks in a familiar chamber (the trained group), or exposure to the chamber alone (the control group). We did not present unpaired tones and shocks to the control group because this paradigm constitutes a different type of associative learning and results in plasticity at LA synapses ^32,43^. Long-term memory formation requires d*e novo* translation during a critical period of several hours after training ^5,31^, thus we sacrificed animals during this time window and collected separate tissue blocks containing either the auditory cortex or the amygdala. Although we refer to these samples as cortex and axons, the cortex samples also contain the proximal axon segments, myelinated segments that pass through the dorsal portion of the external capsule, as well as intrinsic projections and corticocortical projections terminating in adjacent areas of TE1 and perirhinal cortex ^26,27^.

**Figure 3.**
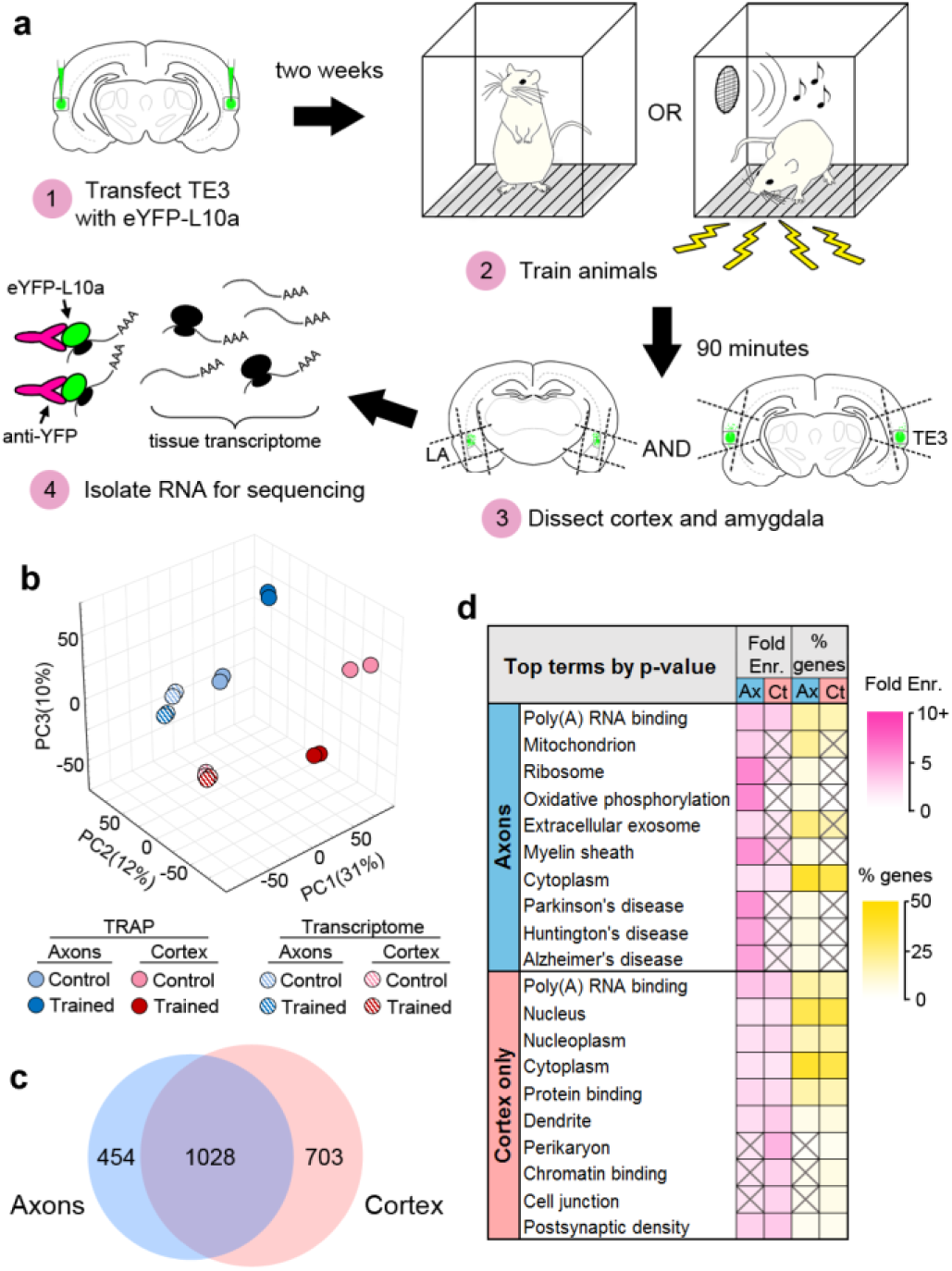
Isolation of the TE3 axonal translatome. a) Experimental workflow (see text). b) Principal component analysis of all experimental replicates. c) Overlap between axonal and cortical translatomes. d) Most enriched GO terms and KEGG pathways in axonal and cortex-only translatomes, sorted by Benjamini- Hochberg adjusted p-value. Gray X’s indicate effects that were not significant (adjusted p-value >0.05).

RNASeq was performed on the TRAPed mRNAs as well as the total mRNA isolated from the homogenized tissue blocks (the tissue transcriptome). Quality control metrics are shown in Supplementary Table 1. Principal component analysis revealed correspondence between experimental replicates, as well as separation between the TRAPed samples and the transcriptome, the cortex and axons, and the trained and control groups (Figure 3b). Gene expression levels were correlated between replicates (Supplementary Figure 2a). Differential gene expression (DGE) analysis was used to identify genes enriched in the translatome versus the tissue transcriptome for each group, as well as genes differentially expressed between the axons and the cortex in each experimental condition and between the experimental groups in each area (Supplementary Table 2). Comparison with a cell-type-specific proteome ^44^ revealed that neuronal genes were more likely than non-neuronal genes to be enriched in the TRAPed samples versus the tissue transcriptome, whereas non-neuronal genes were more likely to be depleted (Supplementary Figure 2b), confirming that our TRAPed samples contain mainly neuronal genes.

Because no translatome or transcriptome of adult forebrain axons has been previously published, we chose to take a conservative approach to identifying axonal genes in our dataset (Supplementary Figure 3a). In order to minimize false positives introduced by the TRAP procedure, only genes that were differentially expressed between TRAPed samples were included. Although this should account for much of the background from the experimental procedures, it does not account for differences between the background transcriptome of the tissue samples, and we therefore excluded genes that were differentially expressed in the corresponding tissue transcriptomes. Finally, genes that were differentially expressed between TRAPed samples were excluded if the enriched sample also was not enriched versus the tissue transcriptome. We defined genes that met these criteria as axonal if they were regulated by learning in the axons, enriched in the axons versus the cortex in either experimental group, or both. Examination of expression levels showed that our filtering method selected for more abundant genes with higher correlation between experimental replicates (Supplementary Figure 3b). Of the 1482 axonal genes identified, the majority (1028) were also either regulated or enriched in the cortex (Figure 3c), and an additional 703 genes were regulated or enriched only in the cortex (defined as “cortex-only” genes).

To directly assess the background introduced by the IP procedure, we repeated the TRAP experiment with a lentivirus encoding eYFP in place of L10a-eYFP. As expected, there was substantial overlap between genes enriched in the TRAP and eYFP-IP samples versus the tissue transcriptome (Supplementary Figure 3c). There were, however, very few learning-regulated mRNAs in the eYFP-IP experiment, and these had little overlap with the TRAPed mRNAs, and even less after the filtering step. Although there was 47% overlap between axonal and cortical genes in the TRAP experiment, there was only 2.5% overlap in the eYFP-IP experiment. These data confirm that the results of our TRAP experiment are not due to background.

### The axonal translatome is diverse

To characterize the axonal translatome, we used DAVID ^45^ (https://david.ncifcrf.gov, version 6.8) to identify Gene Ontology (GO) Terms and KEGG Pathways enriched in the axonal and cortex-only gene sets. Complete results of DAVID analyses are in Supplementary Table 4. The most significantly enriched terms in axons related to mitochondria, translation, and neurodegenerative diseases, whereas cortex-only genes were enriched for terms associated with the cell body, nucleus, and dendrites (Figure 3d). To ensure that our filtering process did not dramatically skew the composition of the final dataset, we also analyzed the unfiltered set of axonal genes. The resulting list of terms was similar, although enrichment levels were lower, consistent with a lower signal-to-noise ratio in the unfiltered data (Supplementary Figure 4a). Comparison between the filtered data from the TRAP and eYFP-IP experiments revealed little similarity between the most enriched GO terms (Supplementary Figure 4b). Manual grouping of significantly enriched terms revealed that terms relating to the presynaptic compartment and cytoskeleton were also predominantly found in axons, along with terms relating to various other cellular functions such as the ubiquitin-proteasome pathway, GTPase signaling, and intracellular transport (Supplementary Figure 5a).

The size and composition of the TE3 axonal translatome are similar to what has been reported in the translatomes of retinal ganglion cell axons^20^ and cortical synaptoneurosomes^10^, the transcriptome of adult hippocampal neuropil^7–9^, and the transcriptomes of axons isolated from cultures of dorsal root ganglion^17,18^, cultured motor neurons^46^, and mixed cortical/hippocampal neurons^19^. We compared these datasets to our axonal and cortex-only translatomes, and found greater overlap with the axonal genes, with 904 of the 1482 genes present in at least one published dataset (Supplementary Figure 5b). Given that these data were obtained from different cell types, preparations, ages, and species, this suggests that at least some aspects of the axonal transcriptome are universal. Interestingly, our axonal translatome had substantially more overlap with datasets from immature versus mature axons, potentially reflecting recapitulation of developmental mechanisms in learning.

### Opposite learning effects in axons and cortex

The majority of genes in the translatome (75%) were regulated by learning, with 19% and 6% of the remainder enriched in the cortex or axons, respectively. 40% of regulated genes showed significant changes in both axons and cortex, and all but one of these (the mitochondrial enzyme *Dlst*) were regulated in opposite directions (Figure 4a). The magnitude of change in the axons and cortex was significantly correlated for these genes, particularly for those downregulated in axons and upregulated in cortex (Figure 4b). Expression levels in the axons and cortex were significantly correlated in both training groups regardless of learning effects, although genes that were upregulated in the axons showed the highest correlation (Supplementary Figure 6a-b). In the control group, genes that were downregulated in axons showed the lowest correlation between the two areas, but this increased in the trained group, particularly for genes that were also upregulated in the cortex. These results suggest that the axonal translatome is not regulated independently, but that compartment-specific translation is coordinated within the cell. This is underscored by the fact that only 63 genes encompassed the 50 most abundant in both areas and conditions (Supplementary Figure 6c). Genes that were upregulated in axons had the highest expression levels in both areas and conditions, further suggesting common regulatory mechanisms (Figure 4c). In contrast to the TRAP experiment, there was no overlap between the 115 genes regulated by learning in axons and the 21 regulated in cortex in the eYFP-IP experiment.

**Figure 4.**
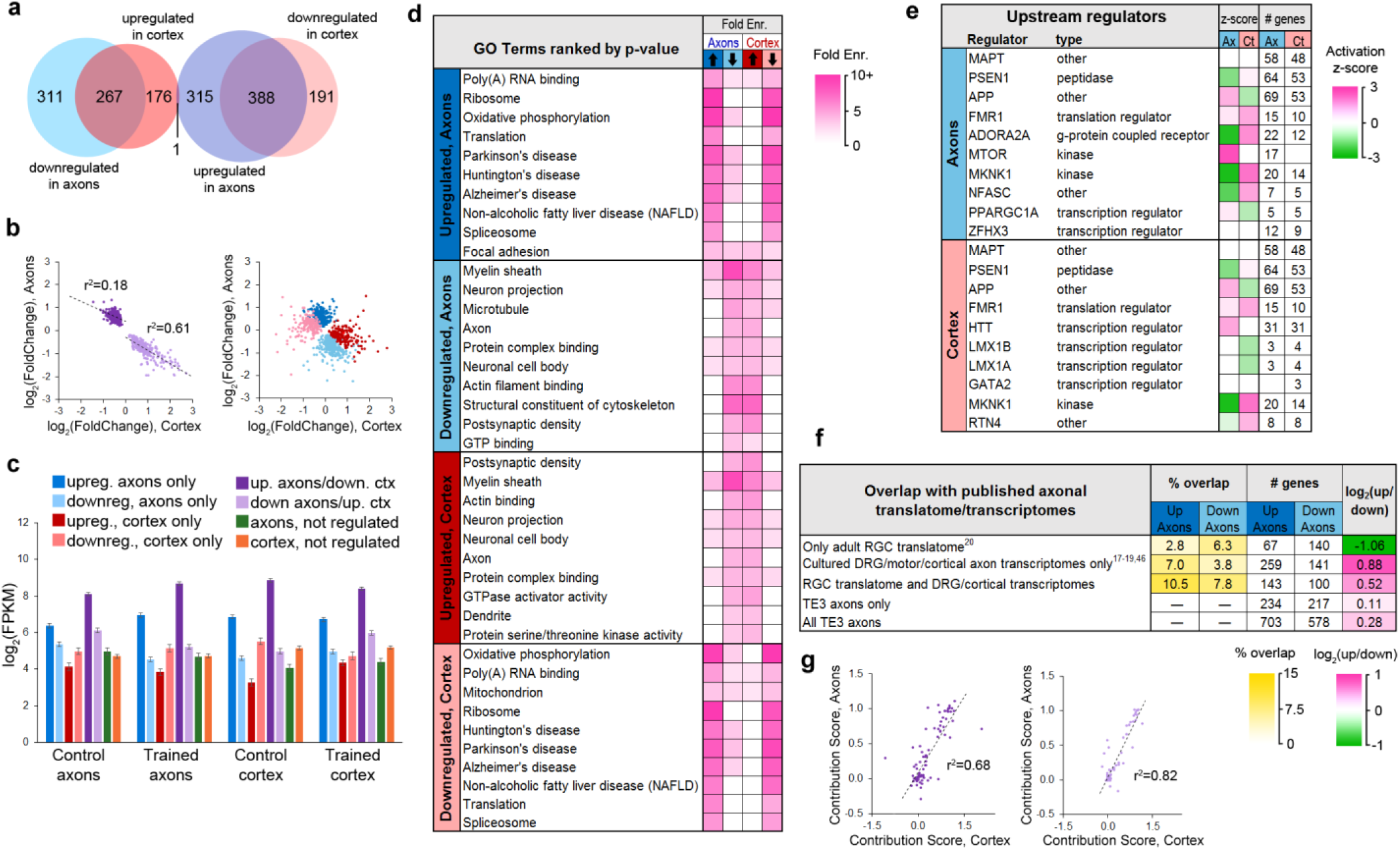
Regulation of the axonal translatome by learning. a) Overlap between learning effects in the axons and cortex. b) Correlations between effect sizes in the axons and cortex for genes differentially expressed in both areas after learning (left) or only one area (right). Regression lines are shown for correlations significant at p<1×10-5. c) Mean expression levels of genes in each group with respect to training effects. Results of ANOVA and post hoc test are given in Supplementary Table 5. Error bars=s.e.m. d) Top GO term and KEGG pathways enriched >3-fold in learning-regulated genes, ranked by Benjamini-Hochberg adjusted p-value. Highly redundant terms are not shown. e) Top regulatory pathways affected by learning in axons and cortex, sorted by adjusted p-value. Activation z-score represents the probability of a pathway being activated or inhibited by learning. f) Overlap between genes up- or downregulated in axons by learning and axonal translatomes and transcriptomes in references 16-19 ^17–20^. g) For genes that had multiple transcripts and were regulated by learning in both axons and cortex, the contribution of each transcript to the gene-level effects in axons and cortex were correlated for genes upregulated in axons and downregulated in cortex (left) and genes downregulated in axons and upregulated in cortex (right). The contribution score was calculated as (change in FPKM transcript)/(change in FPKM gene).

Performing DAVID analysis separately on upregulated and downregulated genes revealed that learning had inverse, function-specific effects on theaxonal and cortical translatomes (Figure 4d). To further explore the effects of learning on cellular functions, we used Ingenuity Pathway Analysis (IPA) software (Qiagen). IPA evaluates changes in gene expression with respect to a database of known pathways and functions, and assigns an enrichment p-value along with a z-score predicting activation or inhibition of a pathway based on published data. A search for upstream regulators found that most of the enriched pathways had opposite z-scores in the axons and cortex (Figure 4e, Supplementary Table 6). Analysis of functional annotations with IPA similarly revealed opposing functional regulation in the two areas (Supplementary Figure 7a, Supplementary Table 7). Although the axonal transcriptome is theoretically a subset of the somatic transcriptome, these results demonstrate an unexpected degree of coordination between the axonal and cortical translatomes.

### Effects of learning on the axonal translatome

Learning affected a range of cellular processes, with some clear patterns of upregulation and downregulation. An overview of regulated genes is shown in Table 1. The genes upregulated in axons, along with those downregulated in cortex, were dominated by two functions: mitochondrial respiration and translation. Axons have high metabolic needs and abundant mitochondria, so it is unsurprising that enrichment of mitochondrial transcripts in axons has been reported by a number of studies^17–20^. Overall, 24% of the transcripts upregulated in axons and 25% of those downregulated in cortex encoded mitochondrial proteins, most of which were involved in either respiration or translation (Figure 4d, Table 1). A few mitochondrial genes were downregulated in axons, however, including some involved in regulation of mitochondrial fusion and localization, such as *Mfn1* and *Opa1*. The opposite pattern was reported in the transcriptome of cultured cortical neurons two days after injury: *Mfn1* was upregulated while transcripts related to respiration were downregulated^19^. If similar regulation occurs in the two paradigms, these results are consistent with translation of dormant axonal mRNAs in response to activity, leading to their upregulation in the translatome and subsequent depletion from the transcriptome.

**Table 1.**
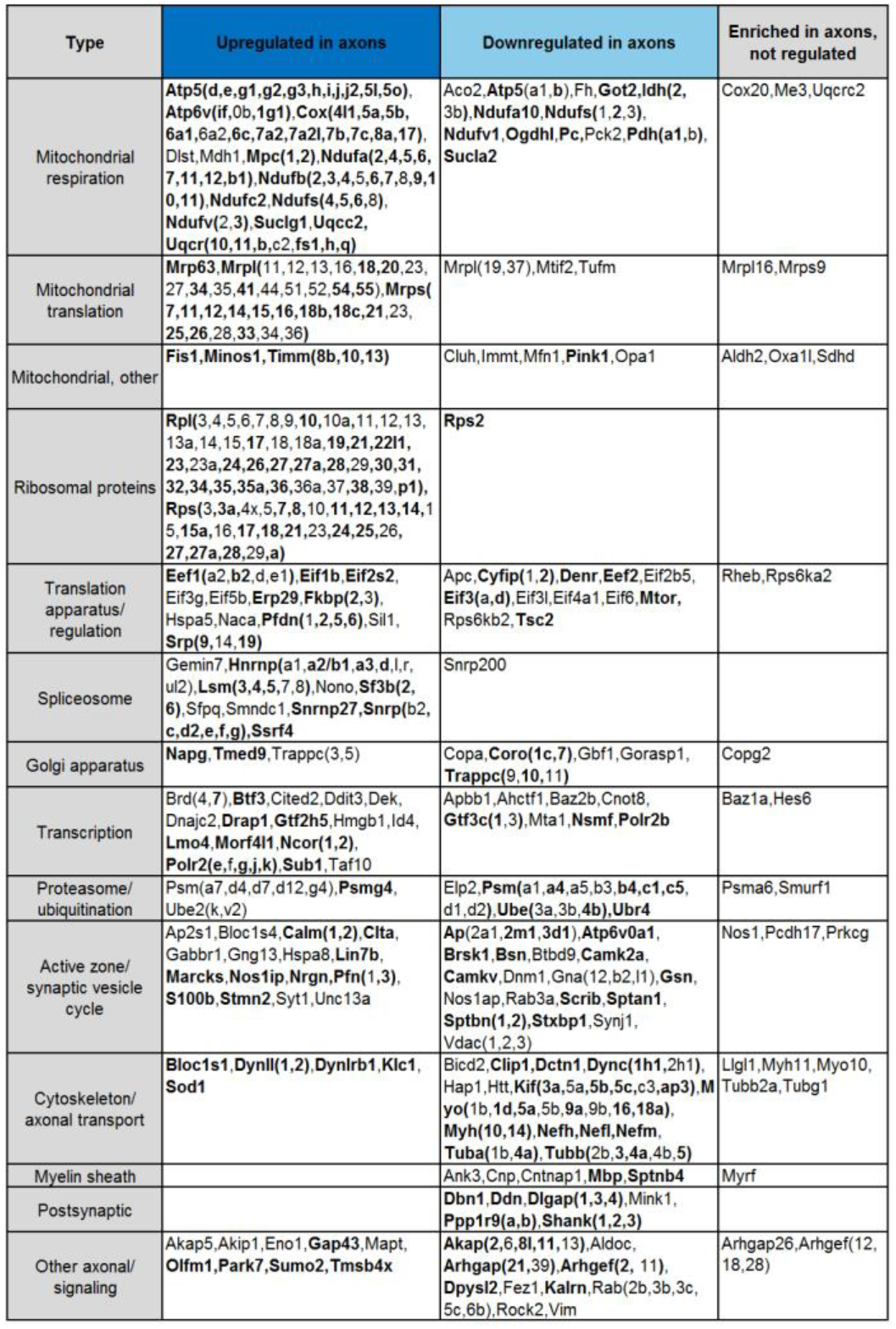
Examples of genes found in auditory cortical axons during memory consolidation by function and effect of learning. Genes in bold type were changed in the opposite direction in the cortex.

Genes coding for translation-related functions, from mRNA splicing to protein folding, were also largely upregulated in axons and downregulated in cortex. Of 68 axonal transcripts encoding ribosomal proteins, 67 were upregulated after learning and 37 of these were downregulated in the cortex. The axonal translatome contained spliceosome components, nearly all of which were upregulated. Genes for initiation and elongation factors were mostly upregulated, although some were downregulated. Intriguingly, a number of genes encoding transcription factors were regulated in axons. Transcription factors are translated locally in growth cones and transported retrogradely to the nucleus (see reference 47 for review), so this could be a case of developmental mechanisms supporting learning in the adult.

A number of transcripts encoding Golgi and rough ER proteins were present in the axonal translatome, although neither of these structures are seen in adult forebrain axons by EM. Similar observations have been reported in axons of cultured neurons, which carry out Golgi and rough ER functions in the absence of classical structures^48–50^. Rough ER proteins tended to be upregulated, whereas Golgi proteins were both upregulated and downregulated. Several upstream regulators of translation were downregulated in axons, including *Apc*, *Cyfip1*, *Mtor*, and *Tsc2*. Because mTOR complex 1 (mTORC1) activates translation of ribosomal proteins and translation factors^51,52^, one possibility is that *Mtor* mRNA was depleted from axons in an initial wave of learning-induced translation, leading to upregulated translation of downstream targets at the time the tissue was collected. Consistent with this, IPA analysis indicated activation of mTOR in the axons (Figure 4e).

Mitochondrial and ribosomal genes made up half of the most highly expressed genes (Supplementary Figure 4c), which could account for the high average expression level of upregulated axonal genes (Figure 4). However, removing these genes did not substantially lower the mean expression levels (Supplementary Figure 6d), indicating that high expression is a feature of upregulated genes independent of function.

Genes downregulated in axons encoded more diverse types of proteins than upregulated genes. These included cytoskeletal components and molecular motors, including tubulins, myosins, dyneins, kinesins, and neurofilaments (Figure 4d, Table 1). Genes encoding synaptic proteins, including synaptic vesicle cycle, active zone, and postsynaptic density proteins, were downregulated, as were signaling molecules and components of the ubiquitin/proteasome pathway and myelin sheath. We used DAVID to examine the 25% of genes in our dataset that were not regulated by learning to determine if there were any functions specific to these genes, but found only one term, “mitochondrion,” enriched in axonal genes, and terms relating to the somatodendritic compartment enriched in the cortex (Supplementary Figure 5a).

We compared the learning-regulated genes to published translatomes of *in vivo* RGC axons^20^ and transcriptomes of cultured DRG and cortical axons^17–19^, and found that genes that overlapped with only the RGC axon translatome were twice as likely to be downregulated as upregulated; in contrast, the converse was true of genes in the cultured axon transcriptomes (Figure 4f). Regulated genes generally had more overlap with datasets from less mature axons, suggesting similar regulation of axonal translation during learning and development (Supplementary Figure 7b). Upregulated genes were much more likely to overlap with genes downregulated rather than upregulated in response to injury^19^, consistent with similar translation patterns leading to depletion from the transcriptome.

To verify axonal localization of mRNA in the amygdala *in vivo*, we used fluorescence *in situ* hybridization (FISH) combined with immunolabeling for axonal neurofilaments. We chose four transcripts that were abundant in control axons and significantly downregulated after learning: the Ras-related protein *Rab3a*, which regulates synaptic vesicle fusion, the N-myc downstream regulated gene *Ndrg4*, the Rab GDP dissociation inhibitor *Gdi1*, and *Ap2m1*, a subunit of the adaptor protein complex 2 which mediates synaptic vesicle endocytosis. Successful FISH labeling required target retrieval treatments, including protease digestion, which proved incompatible with immunolabeling of cytoplasmic GFP in TE3 axons. The monoclonal antibody cocktail SMI 312, which recognizes heavily phosphorylated axonal neurofilaments, was used to identify axons. Rats were given control training and brains were collected at the same time point as in the TRAP experiments. All four mRNA probes, but not the negative control probe, showed punctate labeling in the LA neuropil, with some puncta colocalized with axonal neurofilaments (Figure 5, Supplementary Figure 8).

**Figure 5.**
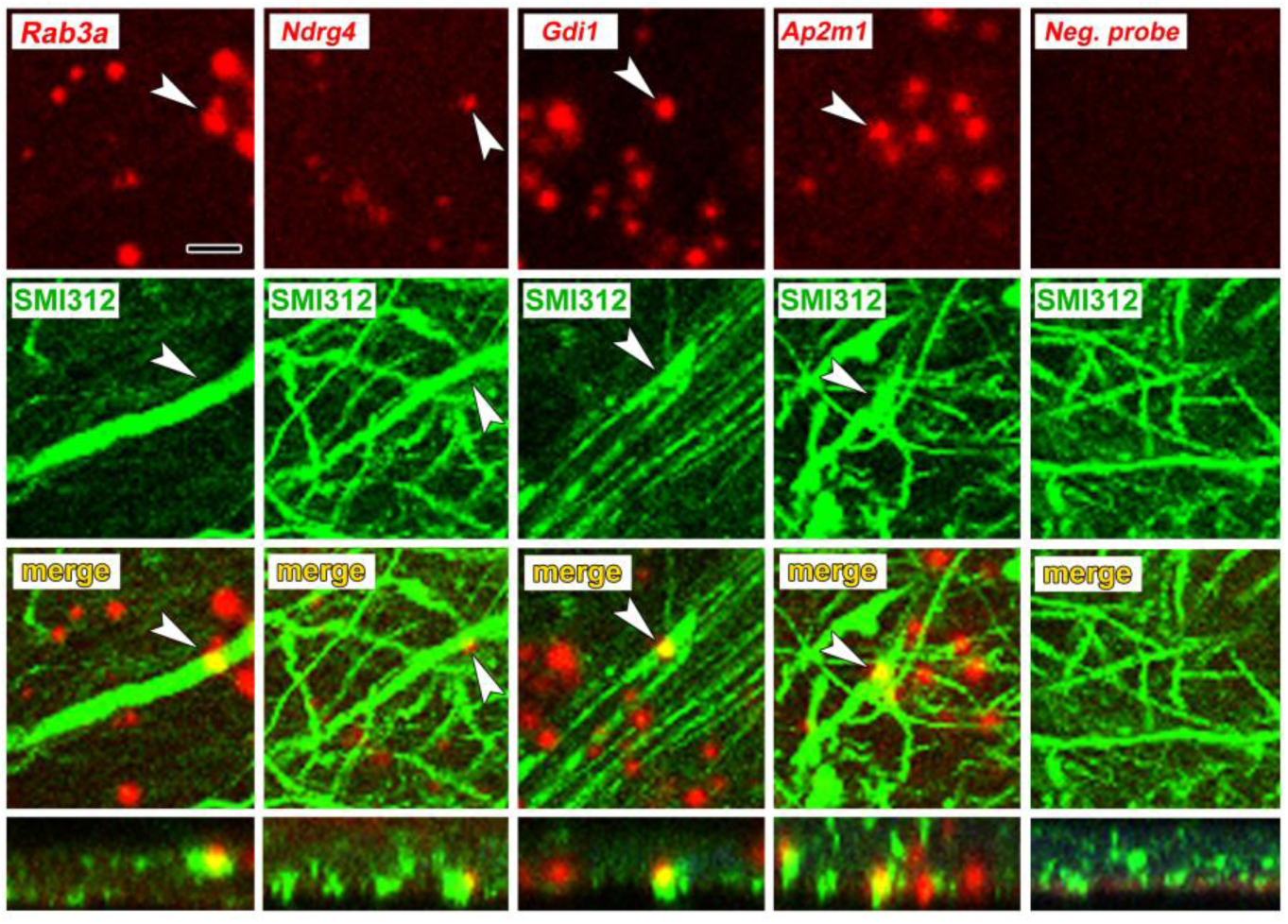
Axonal localization of mRNAs *in vivo*. First row: FISH showing localization of four mRNAs, but not a control probe, in amygdala neuropil. Second and third rows: Immunolabeling with the pan-axonal neurofilament antibody smi-312 shows overlap with mRNA probes. Bottom row: XZ orthogonal view of merged images. Scale = 5 μm.

### Transcript-level correspondence of axonal and cortical mRNA

Because alternative splicing could differ between the axons and cortex, we used Cufflinks software to compare expression at the transcript level (Supplementary Table 9). This analysis identified three genes that were not regulated at the gene level, but had one transcript upregulated (*Gng2)* or downregulated (*Snx27, Speg*) in axons while a second transcript was not (Supplementary Figure 8a). Although multiple transcripts were identified for 133 of the 2185 differentially expressed genes, only one, *Gria2*, had one transcript significantly enriched in axons and another in cortex. Of the 656 genes that were regulated by learning in both the axons and cortex, 54 had more than one transcript, and in 9 cases the same transcript was regulated in both (Supplementary Figure 9b-c). To assess how learning effects were distributed among transcripts in the two areas, we calculated a “contribution score” for each transcript, indicating the fraction of the effect on its parent gene it represents. These scores were correlated between the axons and cortex (Figure 4g), indicating a high degree of coordinated regulation transcript level, similar to that seen at the gene level. Nevertheless, nine genes had transcripts whose axonal and cortical scores differed by >0.3, meaning that more than 30% of the learning effect was on different transcripts (Supplementary Figure 9b-c).

## Discussion

Our results demonstrate that local translation occurs in axons of the adult forebrain *in vivo*, and that the axonal translatome within a memory circuit is regulated by learning. This supports a growing body of evidence that mature axons are capable of local translation, contrary to traditional assumptions, and suggests that gene expression is more extensively decentralized than previously thought. A striking and unexpected feature of our data was the extent of opposing changes in the cortex and axons, suggesting highly coordinated regulation between the two compartments. In dendrites, mRNA transport is activity-regulated, with different trafficking mechanisms exist for different mRNAs ^2,6,53^, and the axonal transcriptome could be similarly regulated. Neurotrophic factors have been shown to induce transport of existing mRNAs from the soma into the axons of cultured DRG neurons, and this is selective for transcripts encoding cytoskeletal proteins^50^. The redistribution of transcripts from the soma to the axons could likewise be due to transport induced by learning. A large range of velocities has been reported for mRNA transport in neural processes^53^, and it is unknown whether mRNA travels from cortical cells to their distal projection fields *in vivo* in the timeframe of our experiment.

Because we analyzed ribosome-bound mRNAs, not the total mRNA in cortical cells, our data reflect not only mRNA localization but translation regulation as well. Presumably, downregulated transcripts reflect termination and subsequent degradation, whereas upregulated transcripts represent new initiation, with or without new transcription. After initiation, ribosomes can be stalled on mRNAs, which are subject to regulated transport and reactivation.^54^ In addition, mRNAs can be transported and stored in a dormant state prior to initiation^53^. Rather than being newly trafficked from the soma, transcripts upregulated in the axons could result from unmasking of preexisting axonal mRNAs, and concomitant depletion from the cortex does not preclude upregulation of new, masked transcripts. Transcripts downregulated in the axons could reflect accelerated elongation in response to learning, or activation of stalled ribosomes, potentially with initiation and subsequent stalling of transcripts in the cortex to replenish the axonal supply. It should be noted that because our cortical samples contained intrinsic and corticocortical axons, it is possible that some of our data derive from asynchronous changes in proximal versus distal axons, potentially due to more rapid trafficking of mRNA from the soma or differential regulation in the proximal axons. We found an assortment of initiation factors and genes coding for them, along with spliceosome components, in axons, making it likely that at least some axonal translation is locally initiated. The presence of genes associated with structures surrounding axons, such as myelin basic protein (*Mbp*), spinophilin (*Ppp1r9b*), dendrin (*Ddn*), and the shank proteins (*Shank1, 2, and 3*), could reflect previously unknown axonal functions of these proteins, as perhaps evidenced by the presence of *Mbp* mRNA in unmyelinated cultured axons^17^. Alternatively, this could result either from trans-endocytosis between dendritic spines and axonal boutons^55^ or exosomal transfer between myelin and the axon shaft^14,34^. Translation regulation in axons is likely to be extensively regulated through multiple mechanisms, the details of which are yet to be fully discovered.

The spatiotemporal uncoupling of translation from transcription has unique implications in the brain, which is itself functionally compartmentalized. The increasing use of gene expression to catalog cells and brain areas, along with genetic targeting of brain circuits, will need to be reexamined if axonal translation is widespread in the adult brain. The idea that translation can be spatially regulated has gradually gained acceptance in a number of contexts, but these continue to be considered exceptional circumstances. Our results counter the longstanding assumption that axonal translation does not occur in the adult brain, and the number and variety of transcripts we identified suggests that spatial regulation could be a fundamental component of translation.

## Methods

### Subjects, surgery, and behavior

All animal procedures were approved by the Animal Care and Use Committees of New York University and the University of Connecticut. Subjects were adult male Sprague-Dawley rats weighing ~300g, housed singly on a 12-hour light/dark cycle with *ad libitum* food and water. All procedures were performed during the rats’ light cycle. For virus injections, rats were anesthetized with ketamine/xylazine and given bilateral stereotaxic injections of either 0.2μl AAV-CMV.eYFP-L10a or 1μl lenti-CMV.eYFP-L10a or lenti-CMV.eYFP (Emory Neuroscience Viral Vector Core) into TE3 (AP 3.8, ML 6.8, DV 3.7mm from interaural center) using a Hamilton syringe. Animals were given at least two weeks to recover from surgery before experiments began.

Behavioral training took place in a soundproof, lit 28.5 x 26 x 28.5cm chamber (Coulbourn Instruments). Auditory tones (30s, 5kHz, 80dB) were delivered through a speaker inside the chamber, and footshocks (0.7mA, 1s) were delivered through a grid floor. Rats were habituated to the conditioning chamber for 30 minutes for two days prior to training. The training protocol consisted of five tones co-terminating with foot shocks delivered over 32.5 minutes with a variable interval between tone-shock pairings.

### Immunolabeling and electron microscopy

Rats were deeply anesthetized with chloral hydrate (1.5mg/kg) and perfused transcardially with 500 ml of mixed aldehydes at pH 7.4 at a rate of 75ml/minute with a peristaltic pump. For eYFP immunolabeling, two lentivirus-injected and two uninjected rats were perfused with 0.25% glutaraldehyde/4% paraformaldehyde/4mM MgCl_2_/2mM CaCl_2_ in 0.1M PIPES buffer. For eIF4E and eIF4G labeling six rats were perfused with 0.5% glutaraldehyde/4% paraformaldehyde/4mM MgCl_2_/2mM CaCl_2_ in 0.1M PIPES buffer and alternate sections were used for each antibody. For eIF2α six rats were perfused with 0.25% glutaraldehyde/4% paraformaldehyde in 0.1M phosphate buffer. For ribosomal protein S6 labeling, three rats were perfused with 0.1% glutaraldehyde/4% paraformaldehyde in 0.1M phosphate buffer. Aldehydes and PIPES buffer were obtained from Electron Microscopy Sciences, phosphate buffer and salts were from Sigma-Aldrich. Brains were removed and immersed in the perfusion fixative for one hour before rinsing in buffered saline (0.01M fixation buffer with 154 mM NaCl) and sectioning at 40μm on a vibrating slicer. Sections were blocked for 15 minutes in 0.1% sodium borohydride, rinsed in buffer, and blocked in 1% bovine serum albumin (BSA; Jackson Labs) before overnight incubation in primary antibody in 1% BSA at room temperature. Sections were rinsed, incubated in 1:200 biotinylated goat anti-rabbit or goat anti-mouse (Vector Labs) in 1% BSA for 30 minutes, rinsed, incubated in avidin/biotin complex peroxidase reagent (Vector Labs Vectastain Elite ABC PK-6100) for 30 minutes, then reacted 5 minutes with 1mM 3,3 diaminobenzidine in 0.0015% H_2_O_2_.

All sections from the brains injected with LV-CMV-eYFP-L10a were examined to confirm that there were no infected cell bodies outside of the TE3 injection site. The area around the LA was dissected out of the immunolabeled sections for electron microscopy. Tissue was processed for electron microscopy as previously described^32^. Briefly, tissue was postfixed in reduced osmium (1% osmium tetroxide/1.5% potassium ferrocyanide) followed by 1% osmium tetroxide, dehydrated in a graded series of ethanol with 1.5% uranyl acetate, infiltrated with LX-112 resin in acetone, embedded in resin, and cured at 60° for 48 hours. 45nm sections were cut on an ultramicrotome (Leica) and imaged on a JEOL 1200EX-II electron microscope at 25,000X on an AMT digital camera. Images were cropped and contrast adjusted using Photoshop (Adobe).

For quantification of eIF4E immunolabel, serial 45 nm sections (average 97+/-5) were imaged from each of the six samples. A 4 x 4 µm square was defined in the middle of the central section of each series, and every profile within the square was followed through serial sections to determine its identity and whether it contained label within the series. If a profile could not be definitively identified as an axon, dendrite, spine, or glial process within the series, it was classified as unidentified.

### Antibodies

Antibody sources and dilutions for immunohistochemistry were as follows: anti-eIF4E rabbit polyclonal (Bethyl Labs A301-154A, lot# A301-154A-1) 1:500, anti-eIF4G1 mouse polyclonal (Abnova H00001981-A01, lot# 08213-2A9) 1:500, anti-eIF2α mouse monoclonal (Cell Signaling L57A5, lot# 3) 1:500, anti-GFP mouse monoclonal (Invitrogen A11120, clone# 3E6) 1:1000, and anti-neurofilament (highly phosphorylated medium and heavy) mouse monoclonal cocktail (BioLegend SMI 312 Lot# B263754). To confirm antigen recognition by the polyclonals to eIF4E and eIF4G, the primary antibodies were preadsorbed before use with a 10-fold excess of the immunizing peptide obtained from the antibody supplier, which reduced the density of labeled structures by 97-98%. To control for specificity of the GFP antibodies, tissue from animals without viral injections was run in parallel and did not result in labeled structures. For immunoprecipitation of eYFP-L10a, two mouse monoclonal anti-GFP antibodies (HtzGFP-19F7 lot# 1/BXC_4789/0513 and HtzGFP-19C8 lot# 1/BXC_4788/0513; available from the Memorial Sloan-Kettering Cancer Center Monoclonal Antibody Core Facility, New York, NY) were used as described below. SMI 312 is a cocktail of affinity-purified mouse monoclonal antibodies that recognize highly phosphorylated medium and heavy neurofilament polypeptides

### Cloning and virus packaging

pAAV-CMV-eYFP-L10a was a generous gift from Dr. Thomas Launey (RIKEN Brain Science Institute, Wako, Japan^42^). YFP-L10a was excised from pAAV-CMV-eYFP-L10a using Nhe I and Xho I. The ~1.4 kb band was gel purified (QiaQuick Gel Extraction Kit, Qiagen, Hilden, Germany). pLV-eGFP (purchased from Adgene) was digested with Xba I and Sal I, and the ~6.7 kb band was gel purified. The eYFP-L10a and pLV backbone were then ligated according to the manufacturer's protocol (T4 DNA ligase, ThermoFisher Scientific, Springfield Township, NJ). Virus (VSVG.HIV.SIN.cPPT.CMV.eYFP-L10a) was packaged by The University of Pennsylvania Vector Core. Viral titer was 2.29e09 GC (genome copies)/mL.

### Immunoprecipitation and RNA isolation

Exactly two hours after the start of behavioral training, rats (n=10 per group) were deeply anesthetized with chloral hydrate (1.5mg/kg) and perfused transcardially with 20ml ice cold oxygenated artificial cerebrospinal fluid (ACSF) consisting of 125mM NaCl, 3.3mM KCl, 1.2mM NaH_2_PO_4_, 25mM NaHCO_3_, 0.5mM CaCl_2_, 7mM MgSO_4_, and 15mM glucose with 50µM cycloheximide. Brains were quickly removed, blocked coronally around the amygdala and auditory cortex, and the two hemispheres separated and incubated in the perfusion solution for 4-5 minutes. Each hemisphere was then bisected along the rhinal fissure. The cortex of the dorsal half was peeled away from the underlying hippocampus and the area containing TE3 was dissected out. A smaller block containing the amygdala was dissected from the ventral half by peeling away the ventral hippocampus, trimming off the cortex lateral to the external capsule and trimming away the hypothalamus and medial portion of the striatum. The TE3 and amygdala blocks were quickly frozen in liquid nitrogen and stored at −80°C. Control and trained animals were run in parallel and tissue was collected in the middle of the animals’ light cycle.

The polysome purification and RNA extraction were performed according to published protocols^40,42^. TE3 or amygdala tissues from 5 animals were pooled (resulting in 2 biological replicates per group for sequencing), as pilot experiments found that this yielded sufficient mRNA. Samples were homogenized in 2 ml of ice-cold polysome extraction buffer [10mM HEPES, 150mM KCl, 5mMMgCl2, 0.5mM DTT, 1 minitablet Complete-EDTA free Protease Inhibitor Cocktail (Roche), 100µl RNasin^®^ Ribonuclease Inhibitor (Promega) and 100µl SUPERase In™ RNase inhibitor (Ambion), 100µg/ml cycloheximide] in douncer homogenizer. Homogenates were centrifuged for 10 minutes at 2,000 x g at 4°C. The supernatants were clarified by adding 1% IGEPAL^®^ CA-630 (SigmaAldrich) and 30 mM DHPC (Avanti Polar Lipids) and incubated for 5 minutes on ice. The clarified lysates were centrifuged for 15 minutes at 20,000 x g at 4°C to pellet unsolubilized material, and 100µl of the supernatant fluid was collected for isolation of the tissue transcriptome. The remainder was added to the conjugated beads/antibodies (200µl) and incubated at 4C overnight with gentle agitation. The following day, the beads were collected with magnets for 1 minute on ice, then washed in 1mL 0,35M KCl washing buffer (20mM HEPES, 350mM KCl, 5mMMgCl_2_, 0.5mM DTT, 1% IGEPAL^®^ CA-630, 100ul RNasin^®^ Ribonuclease Inhibitor and 100 µl SUPERase In™ RNase inhibitor, 100ug/ml cycloheximide) and collected with magnets.

The conjugated beads/antibodies were freshly prepared before the homogenization on the day of the experiment by incubating 300 µl of Dynabeads MyOne Streptavidin T1 (ThermoFisher Scientific) with 120µl of 1µg/µl Biotinylated Protein L (ThermoFisher Scientific) for 35 min at room temperature with gentle rotation. Then, the conjugated protein L-beads were washed with 1XPBS and collected with magnets for 3 times. The conjugated protein L-beads were resuspended in 175 µl of 0.15M KCl IP wash buffer (20mM HEPES, 150mM KCl, 5mMMgCl_2_, 0.5mM DTT, 1% IGEPAL^®^ CA-630, 100µl RNasin^®^ Ribonuclease Inhibitor and 100 µl SUPERase In™ RNase inhibitor, 100ug/ml cycloheximide) and incubated for 1h at room temperature with 50µg of each antibody. The beads were then washed 3 times with 0.15M KCl IP wash buffer and resuspended in the same buffer with 30mM DHPC.

The RNA was extracted and purified with Stratagene Absolutely RNA Nanoprep Kit (Agilent Technologies, Santa Clara, CA) according to the manufacturer’s instructions. All the buffers were provided with the kit except otherwise specified. Briefly, the beads were resuspended in Lysis Buffer with ß-mercaptoethanol, incubated for 10 min at room temperature. 80% Sulfolane (Sigma) was added to the samples and the samples were mixed for 5-10sec, then added to an RNA-binding nano-spin cup and washed with a Low Salt Washing Buffer by centrifuge for 1min at 12,000 x g at room temperature. DNA was digested by mixing the DNase Digestion Buffer and the samples for 15 min at 37C. Then, the samples were washed with High Salt Washing Buffer, Low Salt Washing Buffer and centrifuged for 1min at 12,000 x g. Finally, the samples were eluted with Elution Buffer and centrifuge for 5min at 12,000 x g at room temperature. The isolated RNA was stored at −80°C.

### Sequencing and differential gene expression (DGE) analysis

RNASeq libraries were made using the SMART-Seq v4 Ultra Low Input RNA Kit for Illumina Sequencing, with the Low Input Library Prep kit v2(Clontech, Cat # 634890 and 634899, respectively), using 50-200 pg of total RNA. 16 cycles of PCR were used for the cDNA amplification step, and 5 PCR cycles to amplify the library prep. Libraries were run on an Illumina HiSeq 2500 instrument, using a paired end 50 protocol; 8 samples were pooled per lane of a high output paired end flow cell, using Illumina v4 chemistry.

Raw sequencing data were received in FASTQ format. Read mapping was performed using Tophat 2.0.9 against the rn6 rat reference genome. The resulting BAM alignment files were processed using the HTSeq 0.6.1 python framework and respective rn6 GTF gene annotation, obtained from the UCSC database. Subsequently the Bioconductor package DESeq2(3.2) was used to identify differentially expressed genes (DEG). This package provides statistics for determination of DEG using a model based on the negative binomial distribution. The resulting values were then adjusted using the Benjamini and Hochberg’s method for controlling the false discovery rate (FDR). Genes with an adjusted p-value < 0.05 were determined to be differentially expressed. For transcript-level analysis, the Cufflinks suite (version 2.2.1) was used. ANOVAs and *post hoc* Bonferroni tests were run using the STATISTICA software package (StatSoft). Raw sequencing data and analysis are available in the NCBI Gene Expression Omnibus (accession # GSE124592).

### Filtering of DGE results

To isolate the axonal translatome with as few false positives as possible, we employed a stringent filtering strategy to our DGE data. Twelve comparisons were run between the 8 samples: the TRAPed mRNAs from the axons and cortex were compared to each other separately in each of the training conditions, and the conditions were compared to each other separately in the two brain areas. The same analysis was performed on the tissue transcriptome samples, and each of the four TRAPed samples was compared directly to its corresponding transcriptome. To assemble a list of axonal mRNAs, we began with the comparisons between the TRAPed samples, since this should account for much of the IP background. Because of potential background noise and variability between the individual samples preparations, we excluded genes from each TRAP comparison if the same effect was observed in the corresponding transcriptome comparison. In addition, genes enriched in a given comparison between TRAP samples were excluded if they were not also enriched versus the transcriptome. Although both of these steps likely result in many false negatives, particularly among transcripts that are highly abundant or ubiquitous in the tissue, we felt that excluding potential false positives was crucial given the novelty of our dataset.

### Gene Ontology and Ingenuity Pathway Analysis

Gene lists were submitted to the DAVID ^45^ Functional Annotation Chart tool and enrichment data from the GOTERM_BP_DIRECT (biological process), GOTERM_CC_DIRECT (cellular component), and GOTERM_MF_DIRECT (molecular function) gene ontology categories and KEGG_PATHWAY (Kyoto Encyclopedia of Genes and Genomes) category were examined, using a Benjamini-Hochberg adjusted p-value cutoff of <0.05. For comparison of learning effects, all regulated genes in each area were submitted, regardless of any effect or enrichment in the other area.

For Ingenuity Pathway Analysis (Qiagen Bioinformatics), we submitted all genes differentially expressed (adjusted p-value <0.05) between the training groups in the axons and cortex, along with the corrected log_2_(fold change) calculated by DESeq2. We performed a Core Analysis with the reference data restricted to human, mouse and rat genes and nervous system tissue; otherwise the program’s default settings were used.

### Fluorescence *in situ* hybridization

Adult male rats (n=4) were given control training and perfused two hours later with 4% paraformaldehyde in 0.1M phosphate buffer, pH 7.4. Brains were sectioned at 40µm on a vibrating tissue slicer (Leica) and mounted on glass slides. RNA was detected using the RNAscope 2.5 HD RED kit (Advanced Cell Diagnostics, Inc.) according to the manufacturer’s instructions, with the exception that the incubation time for the fifth amplification step was doubled to increase the diameter of the puncta. Each section was labeled with one of five probes: *Rab3a*, *Ndrg4*, *Ap2m1*, *Gdi1*, or *DapB* (negative control). Sections were blocked overnight in 1% bovine serum albumin with 0.1% Triton-X in phosphate buffered saline, then incubated with primary antibody at 1:500 for 48 hours at 4° followed by 1:200 Alexa-488 goat anti-mouse for one hour at room temperature. Slides were stained with DAPI, mounted in Prolong Gold (Invitrogen), and imaged on a Leica TCS SP8 confocal microscope (Leica Microsystems). Z stacks were collected using a 63x 1.40 HC PL APO oil immersion lens and z step size of 0.3 microns. All sections were stained in parallel with the same batches of probes and antibody. Laser intensity and gain were constant for all images and brightness and contrast were not adjusted. Maximum intensity projections were created in ImageJ.

## Endnotes

LO designed the study, LO, ES, RS, and ZD performed experiments, LO, AT, and AH performed analysis, and LO and EK wrote the paper with input from all authors.

## Supplementary Materials

**Table S1.**
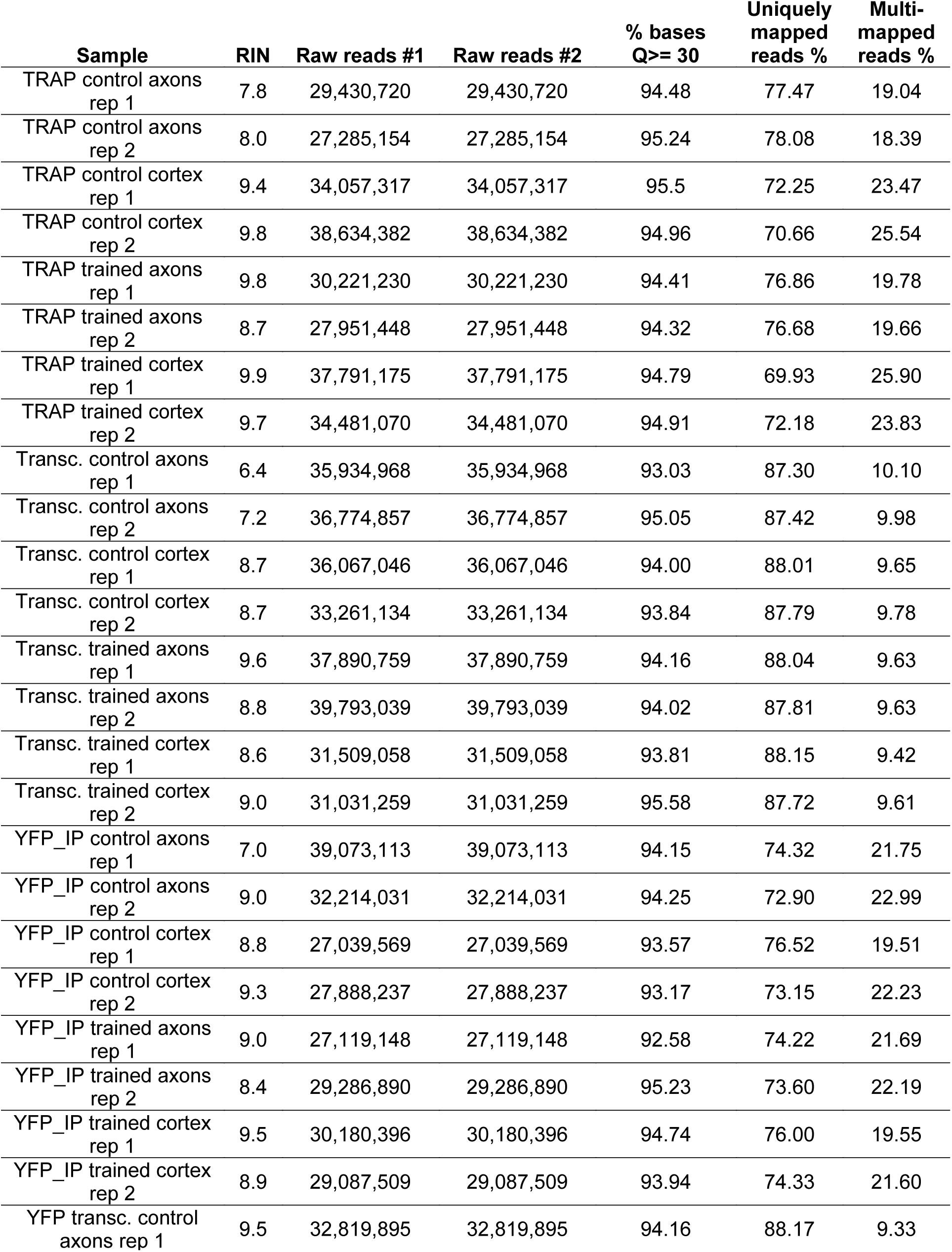
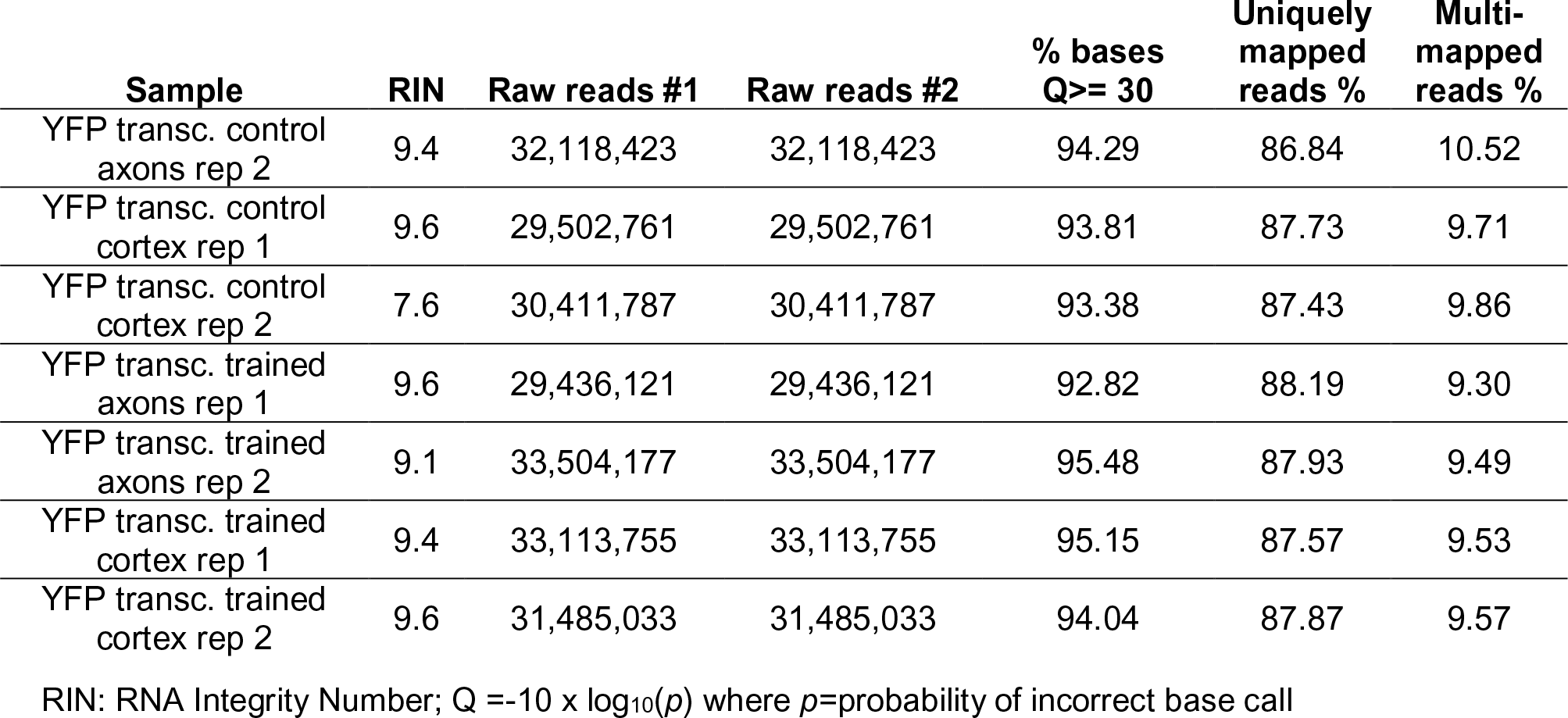
RNA Quality Control Data

**Supplementary Tables 2-8 are in a separate Excel file**

Supplementary Table 2. Results of differential gene expression analysis and subsequent filtering.

Supplementary Table 3. Results of differential gene expression analysis and subsequent filtering, YFP samples.

Supplementary Table 4. Results of DAVID enrichment analyses of all axonal genes, cortex-only genes, and genes that were upregulated and downregulated in the axons and cortex.

Supplementary Table 5. Results of ANOVA and post hoc Bonferroni test comparing mean FPKM between experimental groups by learning effect.

Supplementary Table 6. Results of IPA Upstream Regulator analysis of learning effects in axons and cortex.

Supplementary Table 7. Results of IPA Functional Annotation analysis of learning effects in axons and cortex.

Supplementary Table 8. Transcript-level FPKM values and results of differential expression analysis.

**Supplementary Figure 1.**
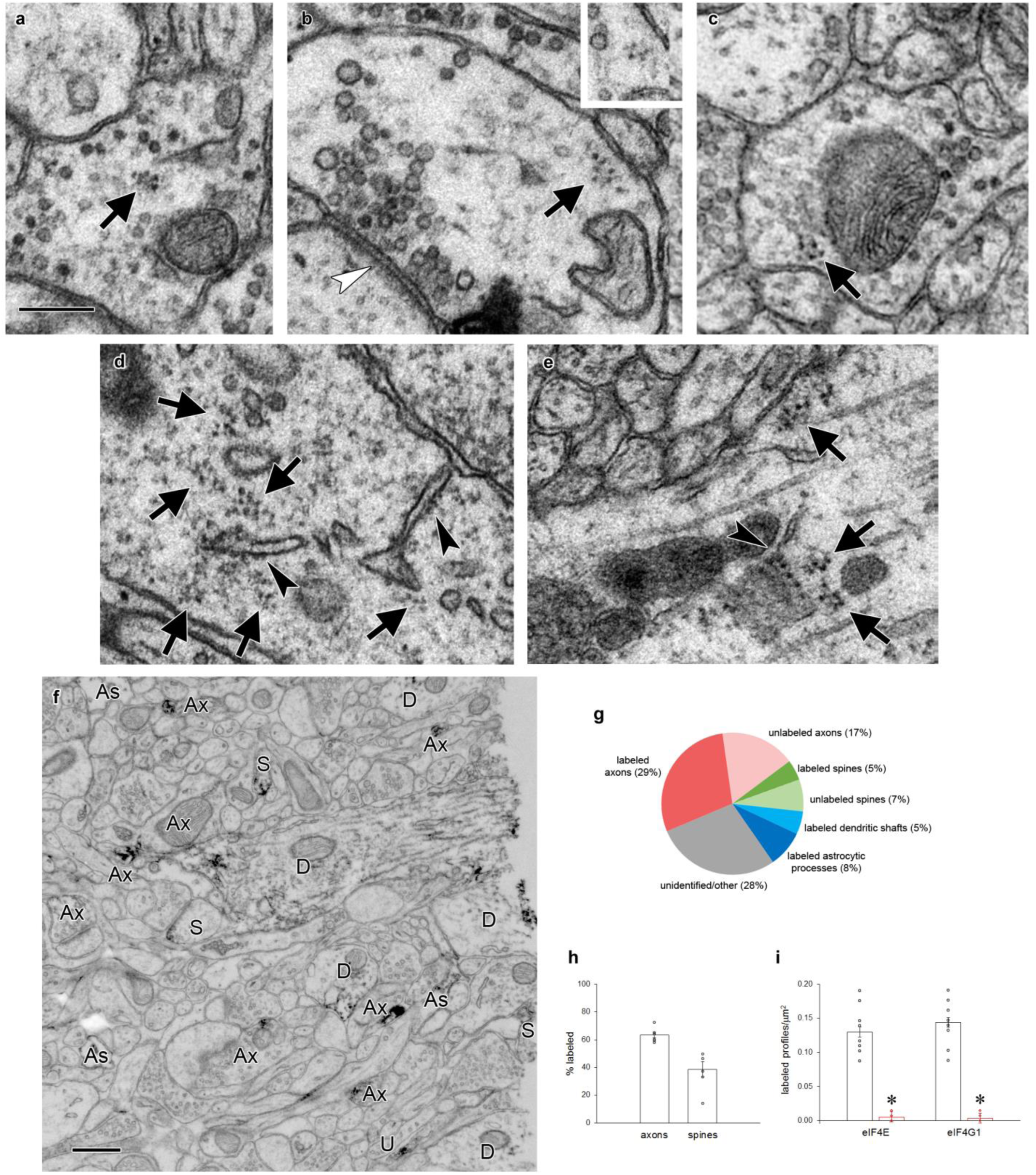
Polyribosomes and translation factors in axons. a-c) Examples of polyribosomes (arrows) in axonal boutons. Inset in (b) shows the same polyribosome on an adjacent serial section. d-e) Copious polyribosomes (arrows) in a neuronal cell body (d) and a large dendritic shaft (e). Rough endoplasmic reticulum (arrowheads) is visible in both structures. f) Representative field of tissue immunolabeled for eIF4E, with labeled axons (Ax), astrocytic processes (As), dendritic shafts (D), and dendritic spines (S) indicated. Profiles were followed through serial sections to confirm identifications. g) Breakdown of all profiles in a 4µm^2^ field of one section near the center of a serial EM volume of tissue immunolabeled for eIF4E. Six series were averaged. 28% of profiles could not be unambiguously identified within the series. h) Percent of axons and spines in a 4µm^2^ field that were immunolabeled for eIF4E when followed through series. 100% of dendritic shafts and astrocytic processes contained label. i) Number of labeled profiles per square micron on 10 randomly chosen, non-consecutive 10 x 10µm electron micrographs of tissue labeled with eIF4E and eIF4G1 antibodies (black) or antibodies preadsorbed with immunizing peptide (red). Densities were compared by ANOVA: eIF4E F_(1,18)_=133.5, p>0.00001; eIF4G1 F_(1,18)_=199.3, p>0.00001. Imaging and analysis were done with experimenters blind to condition.

**Supplementary Figure 2.**
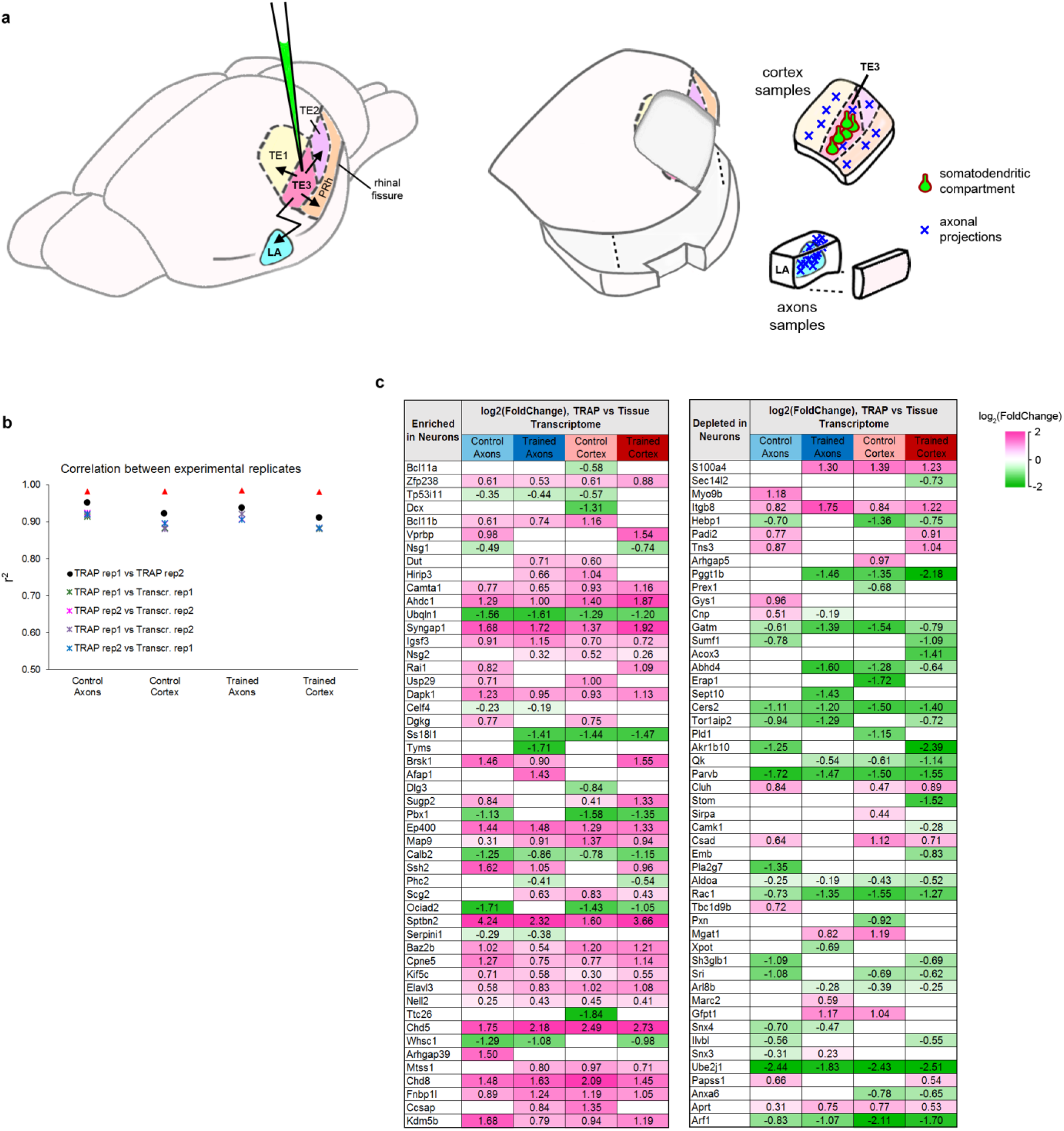
Collection of TRAP samples. a) Left: Illustration of LV-CMV-eYFP-L10a injection into cortical area TE3, showing TE3 projections to cortical areas TE1, TE2, and perirhinal (PRh), and the lateral amygdala (LA). Right: Illustration of tissue sampling for TRAP. After separating the hemispheres and bisecting along the rhinal fissure, cortex samples were collected by dissecting wide margins around TE3 so that portions of adjacent cortical areas and the underlying white matter were included. A separate block was dissected from the ventral half (the “axons” sample), containing the LA, along with the immediately adjacent small area of caudate that also receives projections from TE3. The adjacent area of cortex was removed to ensure that these samples did not contain any stray pieces of perirhinal cortex that could contain cortico-cortical axons. Cortical divisions and projection patterns adapted from references 25-27. b) Correlation coefficients of log_2_(FPKM) between experimental replicates, calculated from all raw data. c) The top genes in the proteome of adult mouse cortex identified as enriched (left) or depleted (right) in neurons versus other cell types, sorted by magnitude of enrichment ^44^. The top 50 genes that were also significantly enriched or depleted in our TRAPed samples versus the tissue transcriptome are shown, with the normalized magnitude of change. Significance was defined as an adjusted p value of <0.05. Neuron-enriched genes were mostly enriched in TRAPed samples (36 of 50), while neuron-depleted genes were depleted from TRAP samples (34 of 50).

**Supplementary Figure 3.**
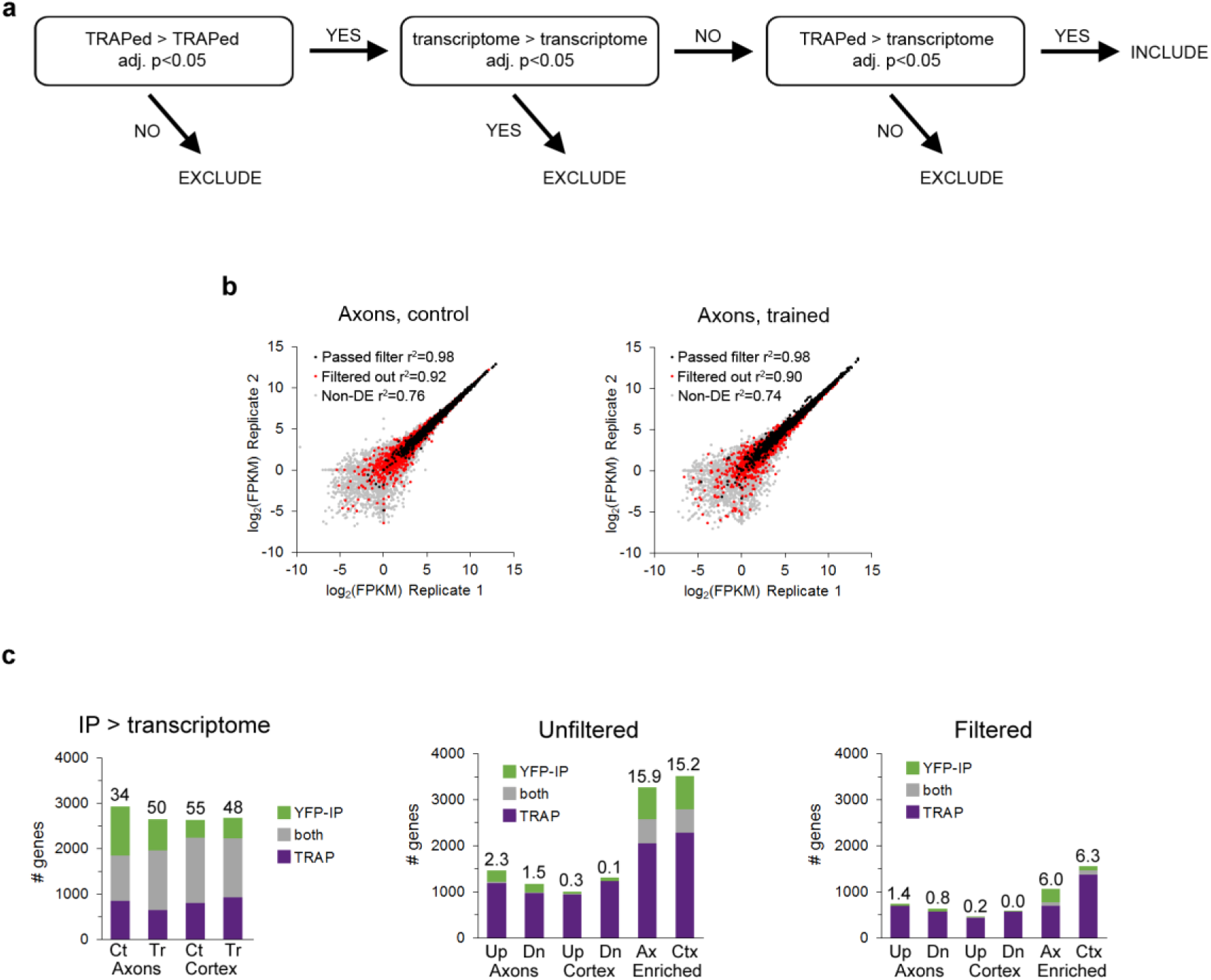
Filtering of DGE results. a) Strategy for removing false positives from results of differential gene expression analysis. b) FPKM values of TRAPed genes from axons in experimental replicates of the control (left) and trained (right) groups. All genes defined as axonal that passed the filtering procedure are indicated with black markers, axonal genes that were removed by filtering with red, and genes that were not axonal in gray. c) Overlap between DGE results in the TRAP and YFP-IP experiments. Left: genes enriched in the TRAP and YFP IP samples versus the transcriptome for all four experimental conditions. Numbers above the bars indicate percent overlap. Center, right: Overlap between genes regulated in axons and cortex (Up, upregulated; Dn, downregulated) or enriched in the axons versus cortex in the unfiltered data (center) and filtered data (right).

**Supplementary Figure 4.**
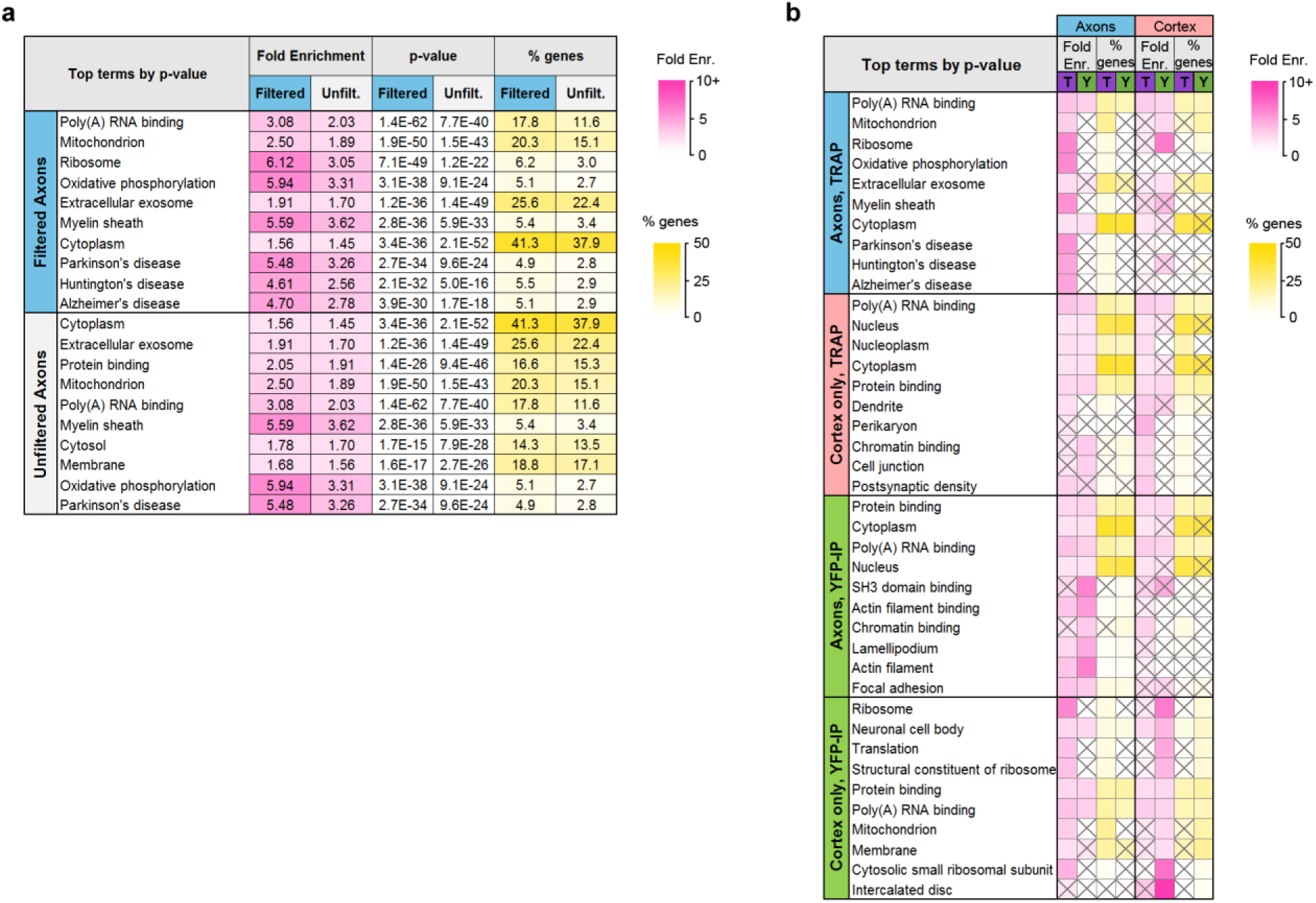
Comparison of TRAP and YFP-IP experiments. a) Top GO and KEGG Pathway terms enriched in the filtered and unfiltered sets of axonal genes, sorted by Benjamini-Hochberg adjusted p-value. b) Top GO Terms and KEGG pathways in axonal and cortex-only translatomes in TRAP and YFP-IP samples, sorted by Benjamini-Hochberg adjusted p-value. Gray X’s indicate effects that were not significant (adjusted p-value >0.05).

**Supplementary Figure 5.**
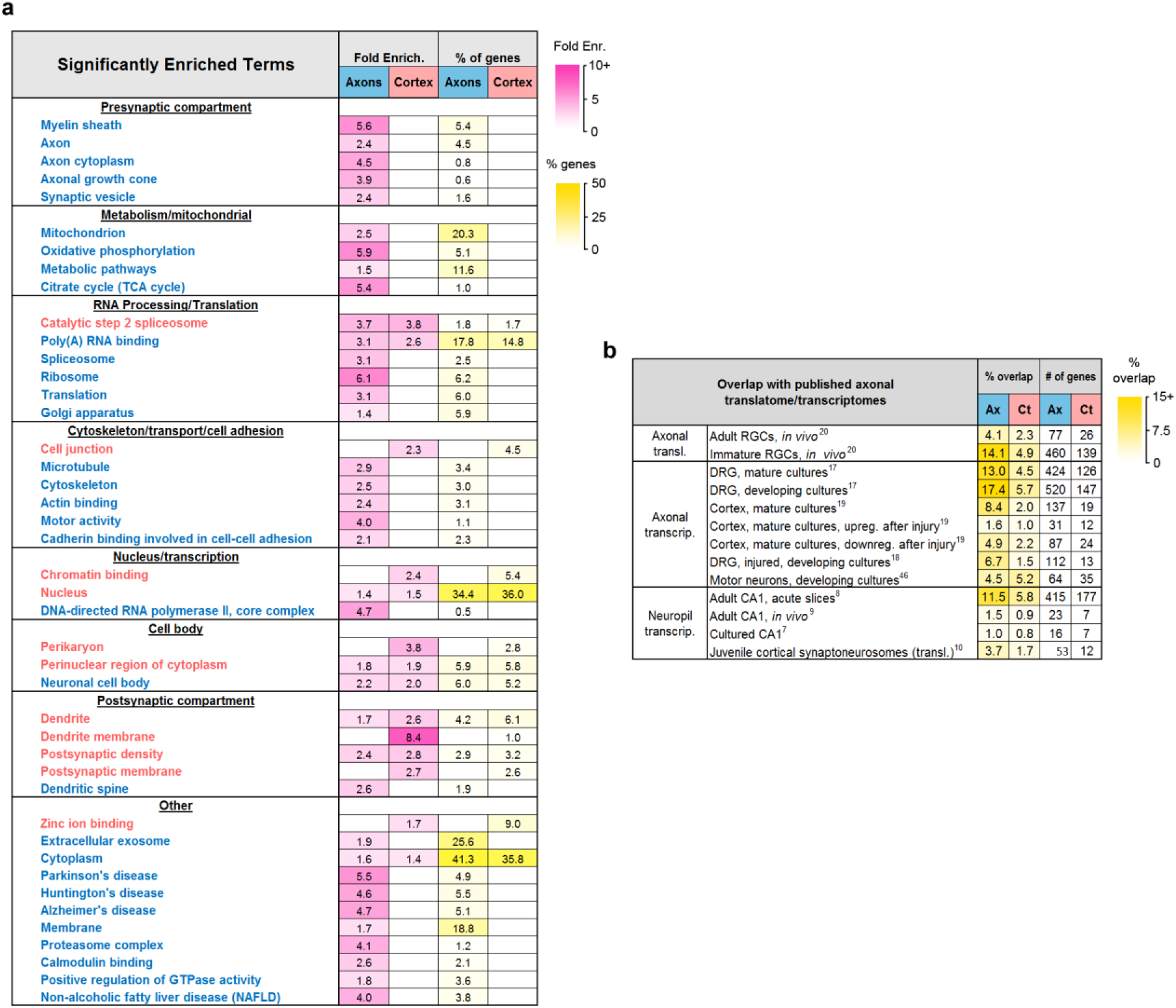
Composition of the axonal translatome. a) Groups of related terms enriched in axonal, cortex-only, or both gene sets. Text color indicates higher enrichment in axons (blue) or cortex (red). Only significant effects (adjusted p-value <0.05) are shown. b) Overlap (% intersection/union) between the axonal and cortex-only and published translatomes and transcriptomes in references 8-10 and 16-19, and number of overlapping genes.

**Supplementary Figure 6.**
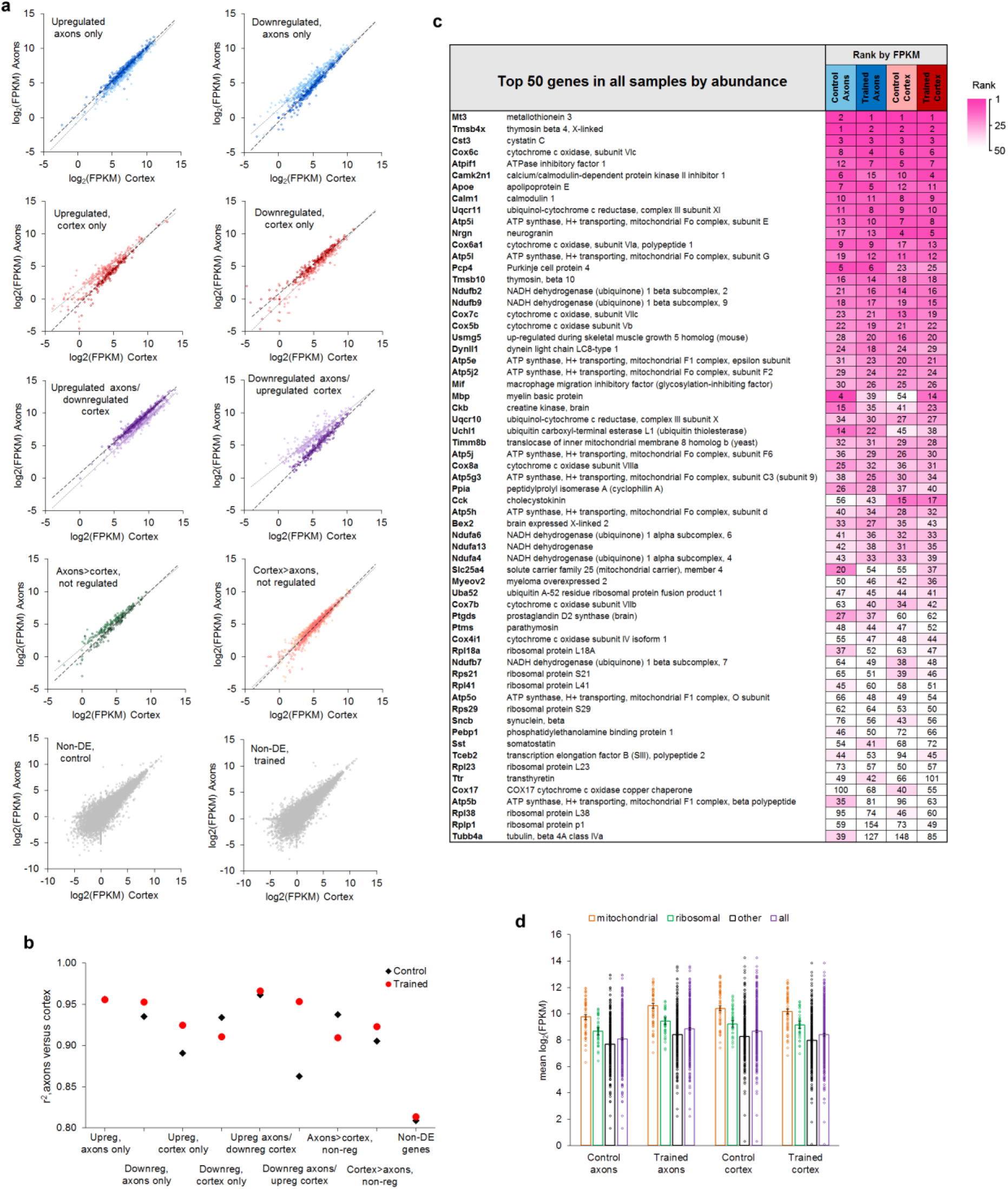
Relative abundance of genes in axons and cortex. a) Plots of log2(FPKM) in cortex versus axons in control (light markers) and trained (dark markers) groups, ^grouped by learning effects. b) Correlation coefficients between log2(FPKM) in cortex and axons for^ each learning effect. c) 63 genes representing the top 50 genes from each of the four groups, sorted by average rank. d) Mean FPKM of genes upregulated in axons and downregulated in cortex after learning, grouped into mitochondrial respiration (n=55), ribosomal proteins (n=39), the remainder (n=294), and the full gene set (n=388). Error bars= s.e.m.

**Supplementary Figure 7.**
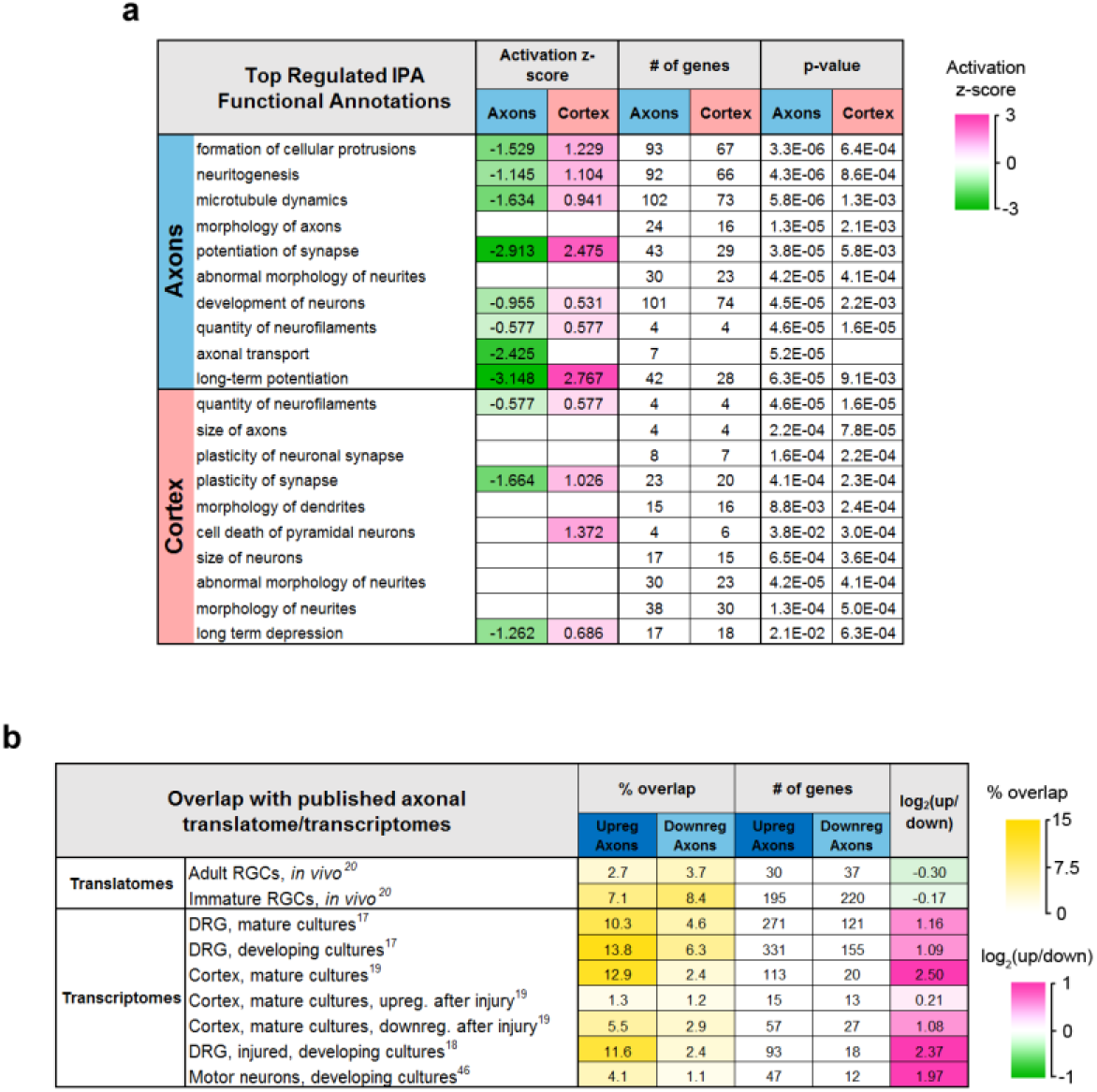
a) Functional annotations significantly regulated by learning in the axons and cortex. b) Overlap between genes regulated in axons and published translatomes and transcriptomes in references (16-19).

**Supplementary Figure 8.**
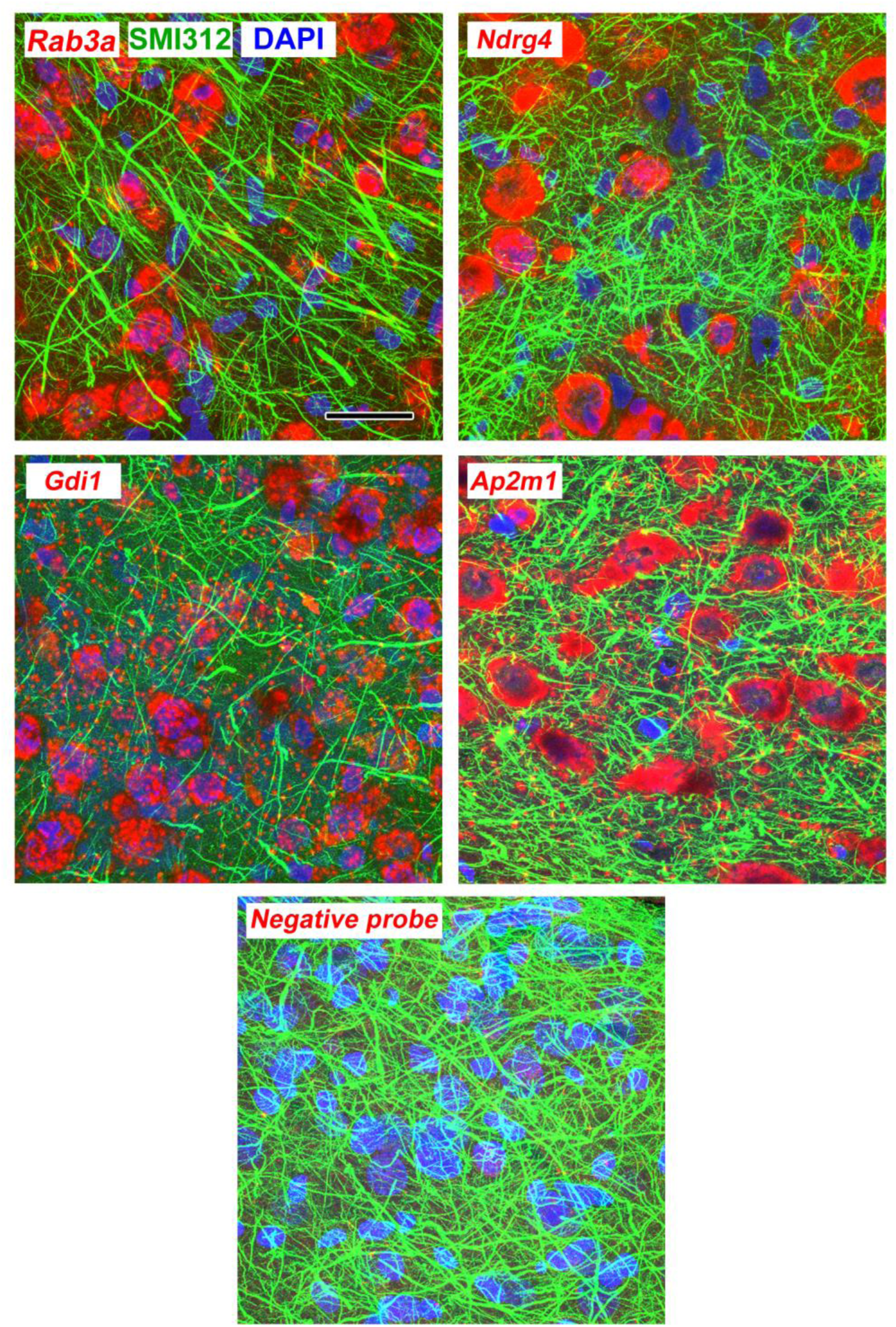
Maximum intensity projections through 3µm (10 confocal images with a 0.3µm z-step size) of lateral amygdala showing FISH labeling and immunolabeling for neurofilaments. Scale = 20 µm.

**Supplementary Figure 9.**
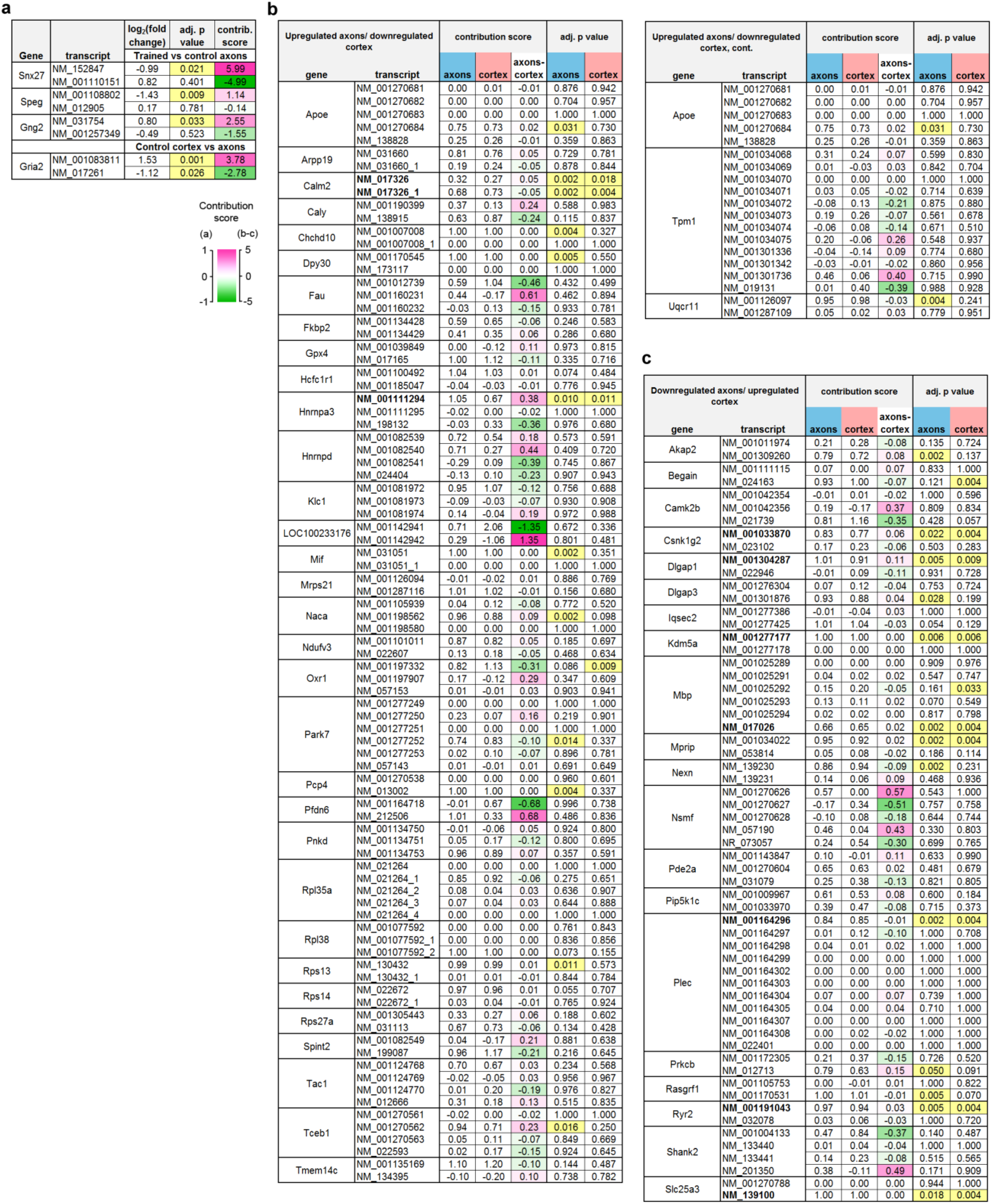
Transcript-level analysis. The contribution score (change in FPKM transcript/change in FPKM gene) indicates the effect of learning on a transcript relative to the net effect on all transcripts of the same gene, with a negative score indicating differences in opposite directions between the transcript and gene. Adjusted p-values for each transcript are highlighted at <0.05. a) Three transcripts were found to be regulated by learning in the axons that were not differentially expressed at the gene level. In each case, a second transcript was affected non-significantly in the opposite direction. The two transcripts of *Gria2* were differently distributed in the control group, with one enriched in axons and the other in cortex. b-c) Genes regulated in both axons and cortex (b; upregulated in axons/downregulated in cortex, c; downregulated in axons/upregulated in cortex) with multiple transcripts in the dataset. The difference between the score in the axons and cortex (“axons – cortex”) indicates the degree of asymmetry, with positive numbers indicating transcripts which were affected proportionally more in the axons than cortex. Values near zero indicate transcripts that were similarly affected in both areas. Transcripts with significant effects in both areas are shown in bold type.

## Acknowledgements

The L10a-YFP construct was a generous gift from Dr. Thomas Launey at the RIKEN Brain Science Institute, Tokyo, Japan. We are grateful to Yutong Zhang for expert technical assistance and to Drs. Joel Richter and Erin Schuman for their insightful comments on the manuscript. This work was supported by NIH grants MH083583 and MH094965 to LO, NS087112 to ES, and NS034007, NS047384, and HD082013 to EK. We would like to thank the Applied Bioinformatics Laboratories (ABL) at the NYU School of Medicine for providing bioinformatics support and helping with the analysis and interpretation of the data. This work has used computing resources at the NYU High Performance Computing Facility (HPCF) and was supported in part by the Viral Vector Core of the Emory Neuroscience NINDS Core Facilities grant, P30NS055077. The Leica SP8 confocal used in this study was obtained with a grant from the NIH (S10OD016435) awarded to Akiko Nishiyama.

