## Supplemental Figures for "The translatome of adult cortical axons is regulated by learning *in vivo*"

### Supplemental Material

#### Table S1. RNA Quality Control Data

| Sample | RIN | Raw reads #1 | Raw reads #2 | % bases Q>= 30 | Uniquely mapped reads % | Multi-mapped reads % |
|---|---|---|---|---|---|---|
| TRAP control axons rep 1 | 7.8 | 29,430,720 | 29,430,720 | 94.48 | 77.47 | 19.04 |
| TRAP control axons rep 2 | 8.0 | 27,285,154 | 27,285,154 | 95.24 | 78.08 | 18.39 |
| TRAP control cortex rep 1 | 9.4 | 34,057,317 | 34,057,317 | 95.5 | 72.25 | 23.47 |
| TRAP control cortex rep 2 | 9.8 | 38,634,382 | 38,634,382 | 94.96 | 70.66 | 25.54 |
| TRAP trained axons rep 1 | 9.8 | 30,221,230 | 30,221,230 | 94.41 | 76.86 | 19.78 |
| TRAP trained axons rep 2 | 8.7 | 27,951,448 | 27,951,448 | 94.32 | 76.68 | 19.66 |
| TRAP trained cortex rep 1 | 9.9 | 37,791,175 | 37,791,175 | 94.79 | 69.93 | 25.90 |
| TRAP trained cortex rep 2 | 9.7 | 34,481,070 | 34,481,070 | 94.91 | 72.18 | 23.83 |
| Transc. control axons rep 1 | 6.4 | 35,934,968 | 35,934,968 | 93.03 | 87.30 | 10.10 |
| Transc. control axons rep 2 | 7.2 | 36,774,857 | 36,774,857 | 95.05 | 87.42 | 9.98 |
| Transc. control cortex rep 1 | 8.7 | 36,067,046 | 36,067,046 | 94.00 | 88.01 | 9.65 |
| Transc. control cortex rep 2 | 8.7 | 33,261,134 | 33,261,134 | 93.84 | 87.79 | 9.78 |
| Transc. trained axons rep 1 | 9.6 | 37,890,759 | 37,890,759 | 94.16 | 88.04 | 9.63 |
| Transc. trained axons rep 2 | 8.8 | 39,793,039 | 39,793,039 | 94.02 | 87.81 | 9.63 |
| Transc. trained cortex rep 1 | 8.6 | 31,509,058 | 31,509,058 | 93.81 | 88.15 | 9.42 |
| Transc. trained cortex rep 2 | 9.0 | 31,031,259 | 31,031,259 | 95.58 | 87.72 | 9.61 |
| YFP_IP control axons rep 1 | 7.0 | 39,073,113 | 39,073,113 | 94.15 | 74.32 | 21.75 |
| YFP_IP control axons rep 2 | 9.0 | 32,214,031 | 32,214,031 | 94.25 | 72.90 | 22.99 |
| YFP_IP control cortex rep 1 | 8.8 | 27,039,569 | 27,039,569 | 93.57 | 76.52 | 19.51 |
| YFP_IP control cortex rep 2 | 9.3 | 27,888,237 | 27,888,237 | 93.17 | 73.15 | 22.23 |
| YFP_IP trained axons rep 1 | 9.0 | 27,119,148 | 27,119,148 | 92.58 | 74.22 | 21.69 |
| YFP_IP trained axons rep 2 | 8.4 | 29,286,890 | 29,286,890 | 95.23 | 73.60 | 22.19 |
| YFP_IP trained cortex rep 1 | 9.5 | 30,180,396 | 30,180,396 | 94.74 | 76.00 | 19.55 |
| YFP_IP trained cortex rep 2 | 8.9 | 29,087,509 | 29,087,509 | 93.94 | 74.33 | 21.60 |
| YFP transc. control axons rep 1 | 9.5 | 32,819,895 | 32,819,895 | 94.16 | 88.17 | 9.33 |

**Table S1. RNA Quality Control Data, cont.**

| Sample | RIN | Raw reads #1 | Raw reads #2 | % bases Q>= 30 | Uniquely mapped reads % | Multi-mapped reads % |
|---|---|---|---|---|---|---|
| YFP transc. control axons rep 2 | 9.4 | 32,118,423 | 32,118,423 | 94.29 | 86.84 | 10.52 |
| YFP transc. control cortex rep 1 | 9.6 | 29,502,761 | 29,502,761 | 93.81 | 87.73 | 9.71 |
| YFP transc. control cortex rep 2 | 7.6 | 30,411,787 | 30,411,787 | 93.38 | 87.43 | 9.86 |
| YFP transc. trained axons rep 1 | 9.6 | 29,436,121 | 29,436,121 | 92.82 | 88.19 | 9.30 |
| YFP transc. trained axons rep 2 | 9.1 | 33,504,177 | 33,504,177 | 95.48 | 87.93 | 9.49 |
| YFP transc. trained cortex rep 1 | 9.4 | 33,113,755 | 33,113,755 | 95.15 | 87.57 | 9.53 |
| YFP transc. trained cortex rep 2 | 9.6 | 31,485,033 | 31,485,033 | 94.04 | 87.87 | 9.57 |

RIN: RNA Integrity Number; Q =-10 x $\log_{10}(p)$ where *p*=probability of incorrect base call

**Supplementary Tables 2-8 are in a separate Excel file**

Supplementary Table 2. Results of differential gene expression analysis and subsequent filtering.

Supplementary Table 3. Results of differential gene expression analysis and subsequent filtering, YFP samples.

Supplementary Table 4. Results of DAVID enrichment analyses of all axonal genes, cortex-only genes, and genes that were upregulated and downregulated in the axons and cortex.

Supplementary Table 5. Results of ANOVA and post hoc Bonferroni test comparing mean FPKM between experimental groups by learning effect.

Supplementary Table 6. Results of IPA Upstream Regulator analysis of learning effects in axons and cortex.

Supplementary Table 7. Results of IPA Functional Annotation analysis of learning effects in axons and cortex.

Supplementary Table 8. Transcript-level FPKM values and results of differential expression analysis.

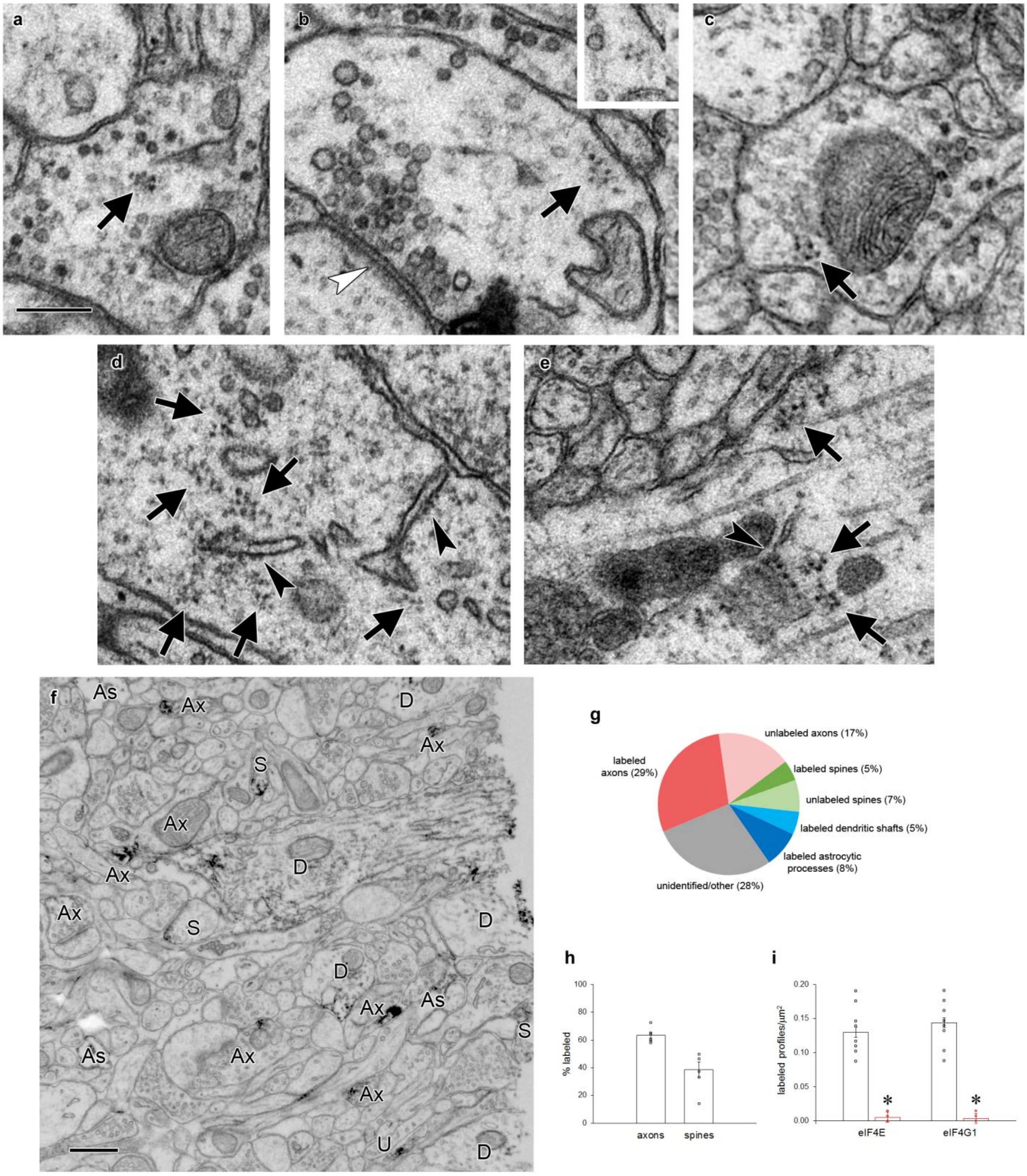
a
b
c
d
e
f
As
Ax
D
S
U
g
unlabeled axons (17%)
labeled axons (29%)
labeled spines (5%)
unlabeled spines (7%)
labeled dendritic shafts (5%)
labeled astrocytic processes (8%)
unidentified/other (28%)
h
% labeled
axons
spines
i
labeled profiles/µm²
eIF4E
eIF4G1
*

**Supplementary Figure 1.** Polyribosomes and translation factors in axons. a-c) Examples of polyribosomes (arrows) in axonal boutons. Inset in (b) shows the same polyribosome on an adjacent serial section. d-e) Copious polyribosomes (arrows) in a neuronal cell body (d) and a large dendritic shaft (e). Rough endoplasmic reticulum (arrowheads) is visible in both structures. f) Representative field of tissue immunolabeled for eIF4E, with labeled axons (Ax), astrocytic processes (As), dendritic shafts (D), and dendritic spines (S) indicated. Profiles were followed through serial sections to confirm identifications. g) Breakdown of all profiles in a $4\mu m^2$ field of one section near the center of a serial EM volume of tissue immunolabeled for eIF4E. Six series were averaged. 28% of profiles could not be unambiguously identified within the series. h) Percent of axons and spines in a $4\mu m^2$ field that were immunolabeled for eIF4E when followed through series. 100% of dendritic shafts and astrocytic processes contained label.

a

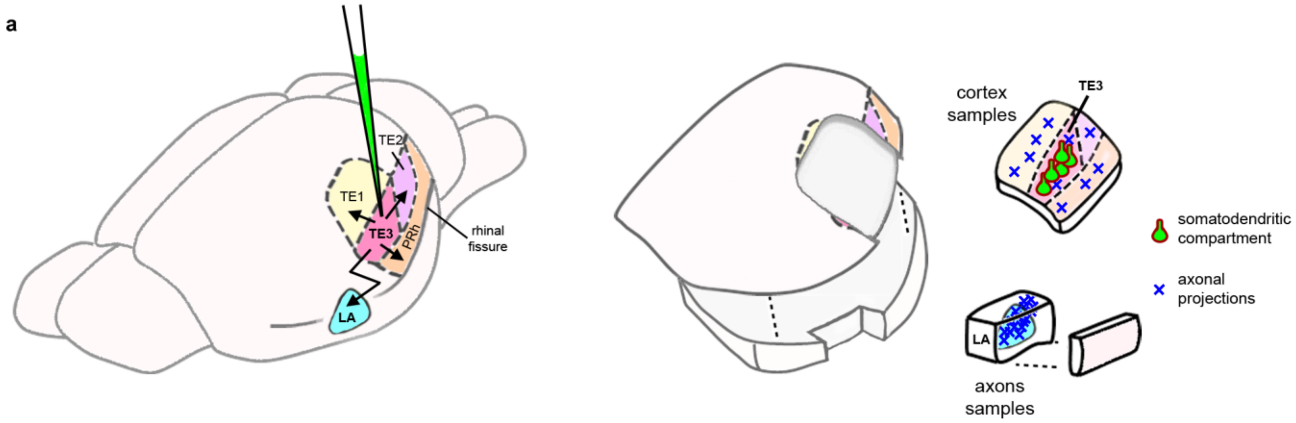


b

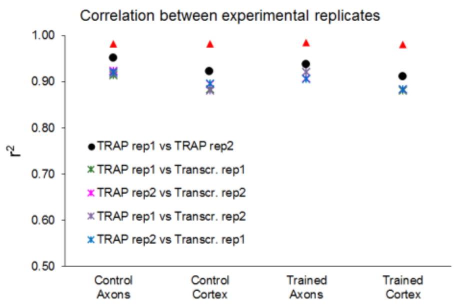


c

| Enriched in Neurons | log2(FoldChange), TRAP vs Tissue Transcriptome | | | |
|---|---|---|---|---|
| | Control Axons | Trained Axons | Control Cortex | Trained Cortex |
| Bcl11a | | | -0.58 | |
| Zfp238 | 0.61 | 0.53 | 0.61 | 0.88 |
| Tp53i11 | -0.35 | -0.44 | -0.57 | |
| Dcx | | | -1.31 | |
| Bcl11b | 0.61 | 0.74 | 1.16 | |
| Vprbp | 0.98 | | | 1.54 |
| Nsg1 | -0.49 | | | -0.74 |
| Dut | | 0.71 | 0.60 | |
| Hirip3 | | 0.66 | 1.04 | |
| Camta1 | 0.77 | 0.65 | 0.93 | 1.16 |
| Ahdc1 | 1.29 | 1.00 | 1.40 | 1.87 |
| Ubqln1 | -1.56 | -1.61 | -1.29 | -1.20 |
| Syngap1 | 1.68 | 1.72 | 1.37 | 1.92 |
| Igsf3 | 0.91 | 1.15 | 0.70 | 0.72 |
| Nsg2 | | 0.32 | 0.52 | 0.26 |
| Rai1 | 0.82 | | | 1.09 |
| Usp29 | 0.71 | | 1.00 | |
| Dapk1 | 1.23 | 0.95 | 0.93 | 1.13 |
| Celf4 | -0.23 | -0.19 | | |
| Dgkg | 0.77 | | 0.75 | |
| Ss18l1 | | -1.41 | -1.44 | -1.47 |
| Tyms | | -1.71 | | |
| Brsk1 | 1.46 | 0.90 | | 1.55 |
| Afap1 | | 1.43 | | |
| Dlg3 | | | -0.84 | |
| Sugp2 | 0.84 | | 0.41 | 1.33 |
| Pbx1 | -1.13 | | -1.58 | -1.35 |
| Ep400 | 1.44 | 1.48 | 1.29 | 1.33 |
| Map9 | 0.31 | 0.91 | 1.37 | 0.94 |
| Calb2 | -1.25 | -0.86 | -0.78 | -1.15 |
| Ssh2 | 1.62 | 1.05 | | 0.96 |
| Phc2 | | -0.41 | | -0.54 |
| Scg2 | | 0.63 | 0.83 | 0.43 |
| Ociad2 | -1.71 | | -1.43 | -1.05 |
| Sptbn2 | 4.24 | 2.32 | 1.60 | 3.66 |
| Serpini1 | -0.29 | -0.38 | | |
| Baz2b | 1.02 | 0.54 | 1.20 | 1.21 |
| Cpne5 | 1.27 | 0.75 | 0.77 | 1.14 |
| Kif5c | 0.71 | 0.58 | 0.30 | 0.55 |
| Elavl3 | 0.58 | 0.83 | 1.02 | 1.08 |
| Nell2 | 0.25 | 0.43 | 0.45 | 0.41 |
| Ttc26 | | | -1.84 | |
| Chd5 | 1.75 | 2.18 | 2.49 | 2.73 |
| Whsc1 | -1.29 | -1.08 | | -0.98 |
| Arhgap39 | 1.50 | | | |
| Mtss1 | | 0.80 | 0.97 | 0.71 |
| Chd8 | 1.48 | 1.63 | 2.09 | 1.45 |
| Fnbp1l | 0.89 | 1.24 | 1.19 | 1.05 |
| Ccsap | | 0.84 | 1.35 | |
| Kdm5b | 1.68 | 0.79 | 0.94 | 1.19 |

| Depleted in Neurons | log2(FoldChange), TRAP vs Tissue Transcriptome | | | |
|---|---|---|---|---|
| | Control Axons | Trained Axons | Control Cortex | Trained Cortex |
| S100a4 | | 1.30 | 1.39 | 1.23 |
| Sec14l2 | | | | -0.73 |
| Myo9b | 1.18 | | | |
| Itgb8 | 0.82 | 1.75 | 0.84 | 1.22 |
| Hebp1 | -0.70 | | -1.36 | -0.75 |
| Padi2 | 0.77 | | | 0.91 |
| Tns3 | 0.87 | | | 1.04 |
| Arhgap5 | | | 0.97 | |
| Pggt1b | | -1.46 | -1.35 | -2.18 |
| Prex1 | | | -0.68 | |
| Gys1 | 0.96 | | | |
| Cnp | 0.51 | -0.19 | | |
| Gatm | -0.61 | -1.39 | -1.54 | -0.79 |
| Sumf1 | -0.78 | | | -1.09 |
| Acox3 | | | | -1.41 |
| Abhd4 | | -1.60 | -1.28 | -0.64 |
| Erap1 | | | -1.72 | |
| Sept10 | | -1.43 | | |
| Cers2 | -1.11 | -1.20 | -1.50 | -1.40 |
| Tor1aip2 | -0.94 | -1.29 | | -0.72 |
| Pld1 | | | -1.15 | |
| Akr1b10 | -1.25 | | | -2.39 |
| Qk | | -0.54 | -0.61 | -1.14 |
| Parvb | -1.72 | -1.47 | -1.50 | -1.55 |
| Cluh | 0.84 | | 0.47 | 0.89 |
| Stom | | | | -1.52 |
| Sirpa | | | 0.44 | |
| Camk1 | | | | -0.28 |
| Csad | 0.64 | | 1.12 | 0.71 |
| Emb | | | | -0.83 |
| Pla2g7 | -1.35 | | | |
| Aldoa | -0.25 | -0.19 | -0.43 | -0.52 |
| Rac1 | -0.73 | -1.35 | -1.55 | -1.27 |
| Tbc1d9b | 0.72 | | | |
| Pxn | | | -0.92 | |
| Mgat1 | | 0.82 | 1.19 | |
| Xpot | | -0.69 | | |
| Sh3glb1 | -1.09 | | | -0.69 |
| Sri | -1.08 | | -0.69 | -0.62 |
| Arl8b | | -0.28 | -0.39 | -0.25 |
| Marc2 | | 0.59 | | |
| Gfpt1 | | 1.17 | 1.04 | |
| Snx4 | -0.70 | -0.47 | | |
| Ilvbl | -0.56 | | | -0.55 |
| Snx3 | -0.31 | 0.23 | | |
| Ube2j1 | -2.44 | -1.83 | -2.43 | -2.51 |
| Papss1 | 0.66 | | | 0.54 |
| Anxa6 | | | -0.78 | -0.65 |
| Aprt | 0.31 | 0.75 | 0.77 | 0.53 |
| Arf1 | -0.83 | -1.07 | -2.11 | -1.70 |

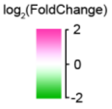


**Supplementary Figure 2.** Collection of TRAP samples. a) Left: Illustration of LV-CMV-eYFP-L10a injection into cortical area TE3, showing TE3 projections to cortical areas TE1, TE2, and perirhinal (PRh), and the lateral amygdala (LA). Right: Illustration of tissue sampling for TRAP. After separating the hemispheres and bisecting along the rhinal fissure, cortex samples were collected by dissecting wide margins around TE3 so that portions of adjacent cortical areas and the underlying white matter were included.  A separate block was dissected from the ventral half (the “axons” sample), containing the LA, along with the immediately adjacent small area of caudate that also receives projections from TE3. The adjacent area of cortex was removed to ensure that these samples did not contain any stray pieces of perirhinal cortex that could contain cortico-cortical axons.  Cortical divisions and projection patterns adapted from references 25-27. b) Correlation coefficients of $\log_2$(FPKM) between experimental replicates, calculated from all raw data. c) The top genes in the proteome of adult mouse cortex identified as enriched (left) or depleted (right) in neurons versus other cell types, sorted by magnitude of enrichment [44]. The top 50 genes that were also significantly enriched or depleted in our TRAPed samples versus the tissue transcriptome are shown, with the normalized magnitude of change. Significance was defined as an adjusted p value of $<0.05$.  Neuron-enriched genes were mostly enriched in TRAPed samples (36 of 50), while neuron-depleted genes were depleted from TRAP samples (34 of 50).

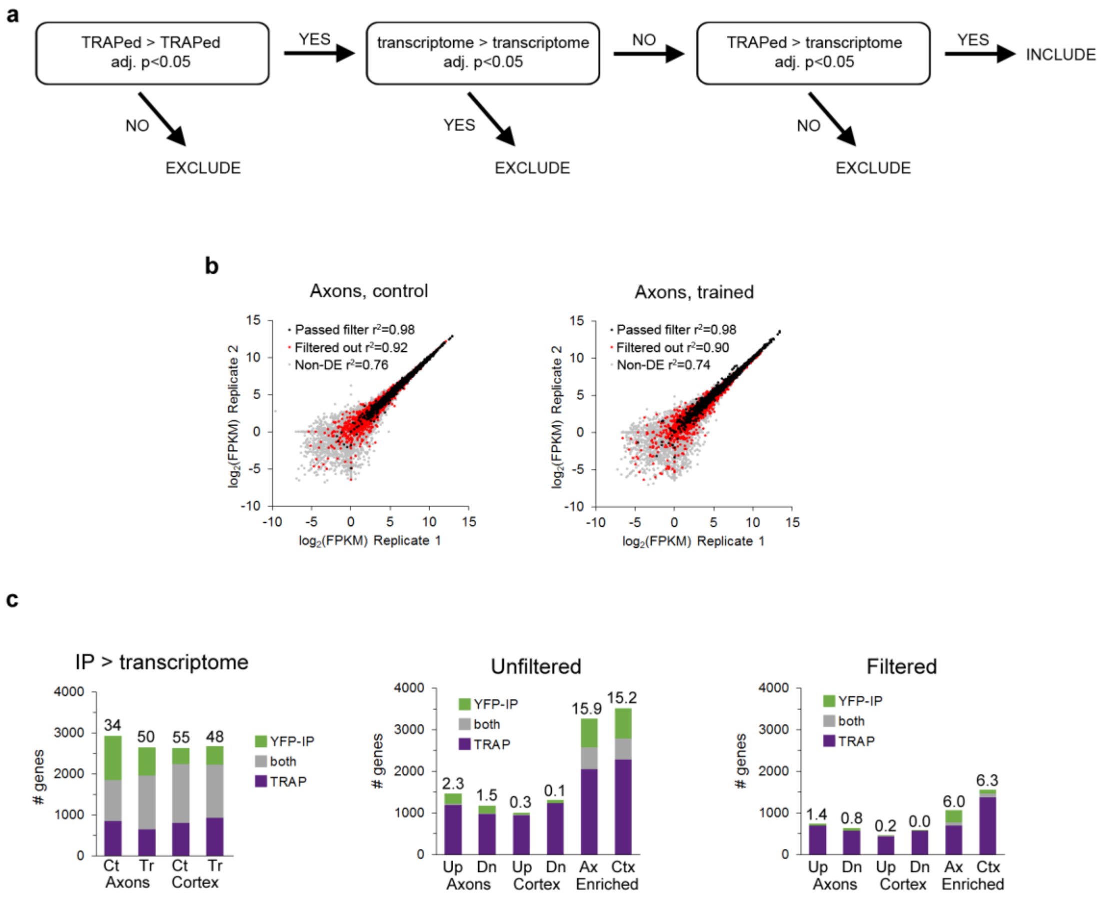


**Supplementary Figure 3.** Filtering of DGE results. a) Strategy for removing false positives from results of differential gene expression analysis. b) FPKM values of TRAPed genes from axons in experimental replicates of the control (left) and trained (right) groups. All genes defined as axonal that passed the filtering procedure are indicated with black markers, axonal genes that were removed by filtering with red, and genes that were not axonal in gray. c) Overlap between DGE results in the TRAP and YFP-IP experiments. Left: genes enriched in the TRAP and YFP IP samples versus the transcriptome for all four experimental conditions. Numbers above the bars indicate percent overlap. Center, right: Overlap between genes regulated in axons and cortex (Up, upregulated; Dn, downregulated) or enriched in the axons versus cortex in the unfiltered data (center) and filtered data (right).

**a**

| | Top terms by p-value | Fold Enrichment | | p-value | | % genes | |
|---|---|---|---|---|---|---|---|
| | | Filtered | Unfilt. | Filtered | Unfilt. | Filtered | Unfilt. |
| Filtered Axons | Poly(A) RNA binding | 3.08 | 2.03 | 1.4E-62 | 7.7E-40 | 17.8 | 11.6 |
| | Mitochondrion | 2.50 | 1.89 | 1.9E-50 | 1.5E-43 | 20.3 | 15.1 |
| | Ribosome | 6.12 | 3.05 | 7.1E-49 | 1.2E-22 | 6.2 | 3.0 |
| | Oxidative phosphorylation | 5.94 | 3.31 | 3.1E-38 | 9.1E-24 | 5.1 | 2.7 |
| | Extracellular exosome | 1.91 | 1.70 | 1.2E-36 | 1.4E-49 | 25.6 | 22.4 |
| | Myelin sheath | 5.59 | 3.62 | 2.8E-36 | 5.9E-33 | 5.4 | 3.4 |
| | Cytoplasm | 1.56 | 1.45 | 3.4E-36 | 2.1E-52 | 41.3 | 37.9 |
| | Parkinson's disease | 5.48 | 3.26 | 2.7E-34 | 9.6E-24 | 4.9 | 2.8 |
| | Huntington's disease | 4.61 | 2.56 | 2.1E-32 | 5.0E-16 | 5.5 | 2.9 |
| | Alzheimer's disease | 4.70 | 2.78 | 3.9E-30 | 1.7E-18 | 5.1 | 2.9 |
| Unfiltered Axons | Cytoplasm | 1.56 | 1.45 | 3.4E-36 | 2.1E-52 | 41.3 | 37.9 |
| | Extracellular exosome | 1.91 | 1.70 | 1.2E-36 | 1.4E-49 | 25.6 | 22.4 |
| | Protein binding | 2.05 | 1.91 | 1.4E-26 | 9.4E-46 | 16.6 | 15.3 |
| | Mitochondrion | 2.50 | 1.89 | 1.9E-50 | 1.5E-43 | 20.3 | 15.1 |
| | Poly(A) RNA binding | 3.08 | 2.03 | 1.4E-62 | 7.7E-40 | 17.8 | 11.6 |
| | Myelin sheath | 5.59 | 3.62 | 2.8E-36 | 5.9E-33 | 5.4 | 3.4 |
| | Cytosol | 1.78 | 1.70 | 1.7E-15 | 7.9E-28 | 14.3 | 13.5 |
| | Membrane | 1.68 | 1.56 | 1.6E-17 | 2.7E-26 | 18.8 | 17.1 |
| | Oxidative phosphorylation | 5.94 | 3.31 | 3.1E-38 | 9.1E-24 | 5.1 | 2.7 |
| | Parkinson's disease | 5.48 | 3.26 | 2.7E-34 | 9.6E-24 | 4.9 | 2.8 |

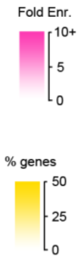


**b**

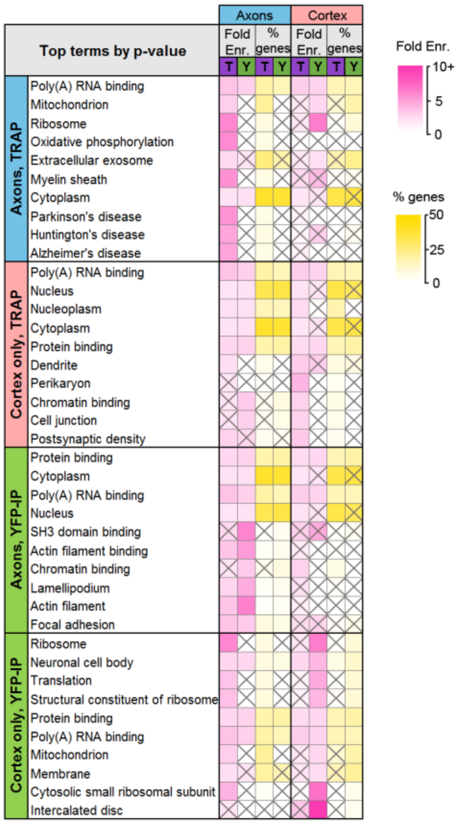


**Supplementary Figure 4.** Comparison of TRAP and YFP-IP experiments. a) Top GO and KEGG Pathway terms enriched in the filtered and unfiltered sets of axonal genes, sorted by Benjamini-Hochberg adjusted p-value. b) Top GO Terms and KEGG pathways in axonal and cortex-only translatomes in TRAP and YFP-IP samples, sorted by Benjamini-Hochberg adjusted p-value. Gray X's indicate effects that were not significant (adjusted p-value >0.05).

**a**

| Significantly Enriched Terms | Fold Enrich. | | % of genes | |
|---|---|---|---|---|
| | Axons | Cortex | Axons | Cortex |
| **Presynaptic compartment** | | | | |
| Myelin sheath | 5.6 | | 5.4 | |
| Axon | 2.4 | | 4.5 | |
| Axon cytoplasm | 4.5 | | 0.8 | |
| Axonal growth cone | 3.9 | | 0.6 | |
| Synaptic vesicle | 2.4 | | 1.6 | |
| **Metabolism/mitochondrial** | | | | |
| Mitochondrion | 2.5 | | 20.3 | |
| Oxidative phosphorylation | 5.9 | | 5.1 | |
| Metabolic pathways | 1.5 | | 11.6 | |
| Citrate cycle (TCA cycle) | 5.4 | | 1.0 | |
| **RNA Processing/Translation** | | | | |
| Catalytic step 2 spliceosome | 3.7 | 3.8 | 1.8 | 1.7 |
| Poly(A) RNA binding | 3.1 | 2.6 | 17.8 | 14.8 |
| Spliceosome | 3.1 | | 2.5 | |
| Ribosome | 6.1 | | 6.2 | |
| Translation | 3.1 | | 6.0 | |
| Golgi apparatus | 1.4 | | 5.9 | |
| **Cytoskeleton/transport/cell adhesion** | | | | |
| Cell junction | | 2.3 | | 4.5 |
| Microtubule | 2.9 | | 3.4 | |
| Cytoskeleton | 2.5 | | 3.0 | |
| Actin binding | 2.4 | | 3.1 | |
| Motor activity | 4.0 | | 1.1 | |
| Cadherin binding involved in cell-cell adhesion | 2.1 | | 2.3 | |
| **Nucleus/transcription** | | | | |
| Chromatin binding | | 2.4 | | 5.4 |
| Nucleus | 1.4 | 1.5 | 34.4 | 36.0 |
| DNA-directed RNA polymerase II, core complex | 4.7 | | 0.5 | |
| **Cell body** | | | | |
| Perikaryon | | 3.8 | | 2.8 |
| Perinuclear region of cytoplasm | 1.8 | 1.9 | 5.9 | 5.8 |
| Neuronal cell body | 2.2 | 2.0 | 6.0 | 5.2 |
| **Postsynaptic compartment** | | | | |
| Dendrite | 1.7 | 2.6 | 4.2 | 6.1 |
| Dendrite membrane | | 8.4 | | 1.0 |
| Postsynaptic density | 2.4 | 2.8 | 2.9 | 3.2 |
| Postsynaptic membrane | | 2.7 | | 2.6 |
| Dendritic spine | 2.6 | | 1.9 | |
| **Other** | | | | |
| Zinc ion binding | | 1.7 | | 9.0 |
| Extracellular exosome | 1.9 | | 25.6 | |
| Cytoplasm | 1.6 | 1.4 | 41.3 | 35.8 |
| Parkinson's disease | 5.5 | | 4.9 | |
| Huntington's disease | 4.6 | | 5.5 | |
| Alzheimer's disease | 4.7 | | 5.1 | |
| Membrane | 1.7 | | 18.8 | |
| Proteasome complex | 4.1 | | 1.2 | |
| Calmodulin binding | 2.6 | | 2.1 | |
| Positive regulation of GTPase activity | 1.8 | | 3.6 | |
| Non-alcoholic fatty liver disease (NAFLD) | 4.0 | | 3.8 | |

Fold Enr.: 10+, 5, 0
% genes: 50, 25, 0

**b**

| Overlap with published axonal translatome/transcriptomes | | % overlap | | # of genes | |
|---|---|---|---|---|---|
| | | Ax | Ct | Ax | Ct |
| Axonal transl. | Adult RGCs, *in vivo*[20] | 4.1 | 2.3 | 77 | 26 |
| | Immature RGCs, *in vivo*[20] | 14.1 | 4.9 | 460 | 139 |
| Axonal transcrip. | DRG, mature cultures[17] | 13.0 | 4.5 | 424 | 126 |
| | DRG, developing cultures[17] | 17.4 | 5.7 | 520 | 147 |
| | Cortex, mature cultures[19] | 8.4 | 2.0 | 137 | 19 |
| | Cortex, mature cultures, upreg. after injury[19] | 1.6 | 1.0 | 31 | 12 |
| | Cortex, mature cultures, downreg. after injury[19] | 4.9 | 2.2 | 87 | 24 |
| | DRG, injured, developing cultures[18] | 6.7 | 1.5 | 112 | 13 |
| | Motor neurons, developing cultures[46] | 4.5 | 5.2 | 64 | 35 |
| Neuropil transcrip. | Adult CA1, acute slices[8] | 11.5 | 5.8 | 415 | 177 |
| | Adult CA1, *in vivo*[9] | 1.5 | 0.9 | 23 | 7 |
| | Cultured CA1[7] | 1.0 | 0.8 | 16 | 7 |
| | Juvenile cortical synaptoneurosomes (transl.)[10] | 3.7 | 1.7 | 53 | 12 |

% overlap: 15+, 7.5, 0

**Supplementary Figure 5.** Composition of the axonal translatome. a) Groups of related terms enriched in axonal, cortex-only, or both gene sets. Text color indicates higher enrichment in axons (blue) or cortex (red). Only significant effects (adjusted p-value <0.05) are shown. b) Overlap (% intersection/union) between the axonal and cortex-only and published translatomes and transcriptomes in references 8-10 and 16-19, and number of overlapping genes.

a

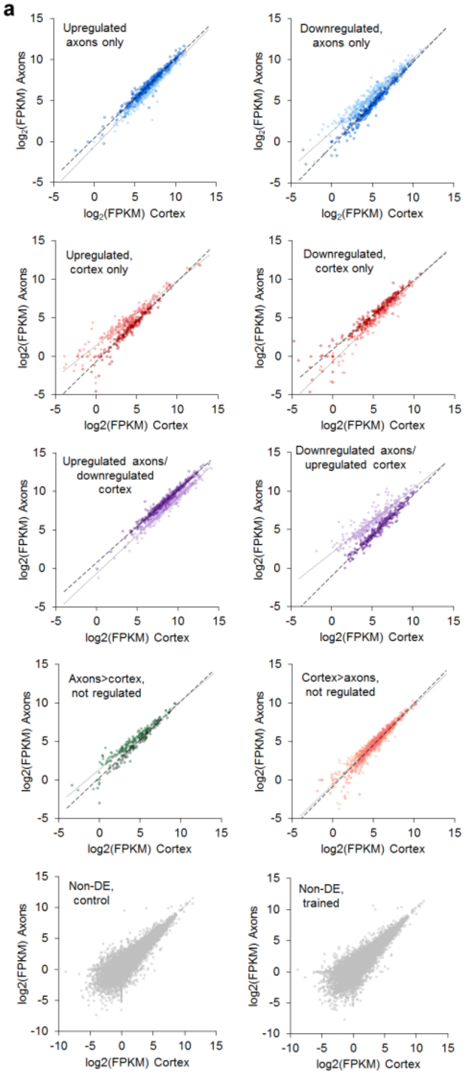


c

| | Top 50 genes in all samples by abundance | Rank by FPKM Control Axons | Trained Axons | Control Cortex | Trained Cortex |
|---|---|---|---|---|---|
| **Mt3** | metallothionein 3 | 2 | 1 | 1 | 1 |
| **Tmsb4x** | thymosin beta 4, X-linked | 1 | 2 | 2 | 2 |
| **Cst3** | cystatin C | 3 | 3 | 3 | 3 |
| **Cox6c** | cytochrome c oxidase, subunit VIc | 8 | 4 | 6 | 6 |
| **Atpif1** | ATPase inhibitory factor 1 | 12 | 7 | 5 | 7 |
| **Camk2n1** | calcium/calmodulin-dependent protein kinase II inhibitor 1 | 6 | 15 | 10 | 4 |
| **Apoe** | apolipoprotein E | 7 | 5 | 12 | 11 |
| **Calm1** | calmodulin 1 | 10 | 11 | 8 | 9 |
| **Uqcr11** | ubiquinol-cytochrome c reductase, complex III subunit XI | 11 | 8 | 9 | 10 |
| **Atp5i** | ATP synthase, H+ transporting, mitochondrial Fo complex, subunit E | 13 | 10 | 7 | 8 |
| **Nrgn** | neurogranin | 17 | 13 | 4 | 5 |
| **Cox6a1** | cytochrome c oxidase, subunit VIa, polypeptide 1 | 9 | 9 | 17 | 13 |
| **Atp5l** | ATP synthase, H+ transporting, mitochondrial Fo complex, subunit G | 19 | 12 | 11 | 12 |
| **Pcp4** | Purkinje cell protein 4 | 5 | 6 | 23 | 25 |
| **Tmsb10** | thymosin, beta 10 | 16 | 14 | 18 | 18 |
| **Ndufb2** | NADH dehydrogenase (ubiquinone) 1 beta subcomplex, 2 | 21 | 16 | 14 | 16 |
| **Ndufb9** | NADH dehydrogenase (ubiquinone) 1 beta subcomplex, 9 | 18 | 17 | 19 | 15 |
| **Cox7c** | cytochrome c oxidase, subunit VIIc | 23 | 21 | 13 | 19 |
| **Cox5b** | cytochrome c oxidase subunit Vb | 22 | 19 | 21 | 22 |
| **Usmg5** | up-regulated during skeletal muscle growth 5 homolog (mouse) | 28 | 20 | 16 | 20 |
| **Dynll1** | dynein light chain LC8-type 1 | 24 | 18 | 24 | 29 |
| **Atp5e** | ATP synthase, H+ transporting, mitochondrial F1 complex, epsilon subunit | 31 | 23 | 20 | 21 |
| **Atp5j2** | ATP synthase, H+ transporting, mitochondrial Fo complex, subunit F2 | 29 | 24 | 22 | 24 |
| **Mif** | macrophage migration inhibitory factor (glycosylation-inhibiting factor) | 30 | 26 | 25 | 26 |
| **Mbp** | myelin basic protein | 4 | 39 | 54 | 14 |
| **Ckb** | creatine kinase, brain | 15 | 35 | 41 | 23 |
| **Uqcr10** | ubiquinol-cytochrome c reductase, complex III subunit X | 34 | 30 | 27 | 27 |
| **Uchl1** | ubiquitin carboxyl-terminal esterase L1 (ubiquitin thiolesterase) | 14 | 22 | 45 | 38 |
| **Timm8b** | translocase of inner mitochondrial membrane 8 homolog b (yeast) | 32 | 31 | 29 | 28 |
| **Atp5j** | ATP synthase, H+ transporting, mitochondrial Fo complex, subunit F6 | 36 | 29 | 26 | 30 |
| **Cox8a** | cytochrome c oxidase subunit VIIIa | 25 | 32 | 36 | 31 |
| **Atp5g3** | ATP synthase, H+ transporting, mitochondrial Fo complex, subunit C3 (subunit 9) | 38 | 25 | 30 | 34 |
| **Ppia** | peptidylprolyl isomerase A (cyclophilin A) | 26 | 28 | 37 | 40 |
| **Cck** | cholecystokinin | 56 | 43 | 15 | 17 |
| **Atp5h** | ATP synthase, H+ transporting, mitochondrial Fo complex, subunit d | 40 | 34 | 28 | 32 |
| **Bex2** | brain expressed X-linked 2 | 33 | 27 | 35 | 43 |
| **Ndufa6** | NADH dehydrogenase (ubiquinone) 1 alpha subcomplex, 6 | 41 | 36 | 32 | 33 |
| **Ndufa13** | NADH dehydrogenase | 42 | 38 | 31 | 35 |
| **Ndufa4** | NADH dehydrogenase (ubiquinone) 1 alpha subcomplex, 4 | 43 | 33 | 33 | 39 |
| **Slc25a4** | solute carrier family 25 (mitochondrial carrier), member 4 | 20 | 54 | 55 | 37 |
| **Myeov2** | myeloma overexpressed 2 | 50 | 46 | 42 | 36 |
| **Uba52** | ubiquitin A-52 residue ribosomal protein fusion product 1 | 47 | 45 | 44 | 41 |
| **Cox7b** | cytochrome c oxidase subunit VIIb | 63 | 40 | 34 | 42 |
| **Ptgds** | prostaglandin D2 synthase (brain) | 27 | 37 | 60 | 62 |
| **Ptms** | parathymosin | 48 | 44 | 47 | 52 |
| **Cox4i1** | cytochrome c oxidase subunit IV isoform 1 | 55 | 47 | 48 | 44 |
| **Rpl18a** | ribosomal protein L18A | 37 | 52 | 63 | 47 |
| **Ndufb7** | NADH dehydrogenase (ubiquinone) 1 beta subcomplex, 7 | 64 | 49 | 38 | 48 |
| **Rps21** | ribosomal protein S21 | 65 | 51 | 39 | 46 |
| **Rpl41** | ribosomal protein L41 | 45 | 60 | 58 | 51 |
| **Atp5o** | ATP synthase, H+ transporting, mitochondrial F1 complex, O subunit | 66 | 48 | 49 | 54 |
| **Rps29** | ribosomal protein S29 | 62 | 64 | 53 | 50 |
| **Sncb** | synuclein, beta | 76 | 56 | 43 | 56 |
| **Pebp1** | phosphatidylethanolamine binding protein 1 | 46 | 50 | 72 | 66 |
| **Sst** | somatostatin | 54 | 41 | 68 | 72 |
| **Tceb2** | transcription elongation factor B (SIII), polypeptide 2 | 44 | 53 | 94 | 45 |
| **Rpl23** | ribosomal protein L23 | 73 | 57 | 50 | 57 |
| **Ttr** | transthyretin | 49 | 42 | 66 | 101 |
| **Cox17** | COX17 cytochrome c oxidase copper chaperone | 100 | 68 | 40 | 55 |
| **Atp5b** | ATP synthase, H+ transporting, mitochondrial F1 complex, beta polypeptide | 35 | 81 | 96 | 63 |
| **Rpl38** | ribosomal protein L38 | 95 | 74 | 46 | 60 |
| **Rplp1** | ribosomal protein p1 | 59 | 154 | 73 | 49 |
| **Tubb4a** | tubulin, beta 4A class IVa | 39 | 127 | 148 | 85 |

b

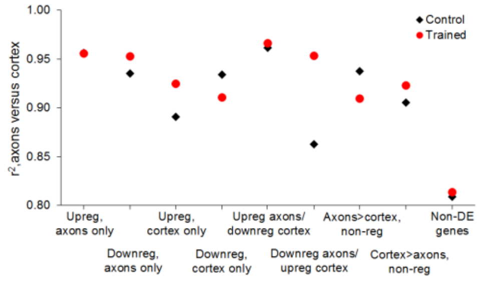


d

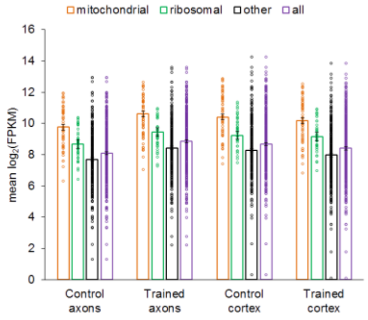


**Supplementary Figure 6.** Relative abundance of genes in axons and cortex. a) Plots of log2(FPKM) in cortex versus axons in control (light markers) and trained (dark markers) groups, grouped by learning effects. b) Correlation coefficients between $\log_2$(FPKM) in cortex and axons for each learning effect. c) 63 genes representing the top 50 genes from each of the four groups, sorted by average rank. d) Mean FPKM of genes upregulated in axons and downregulated in cortex after learning, grouped into mitochondrial respiration (n=55), ribosomal proteins (n=39), the remainder (n=294), and the full gene set (n=388). Error bars= s.e.m.

**a**

| Top Regulated IPA Functional Annotations | | Activation z-score | | # of genes | | p-value | |
|---|---|---|---|---|---|---|---|
| | | Axons | Cortex | Axons | Cortex | Axons | Cortex |
| Axons | formation of cellular protrusions | -1.529 | 1.229 | 93 | 67 | 3.3E-06 | 6.4E-04 |
| | neuritogenesis | -1.145 | 1.104 | 92 | 66 | 4.3E-06 | 8.6E-04 |
| | microtubule dynamics | -1.634 | 0.941 | 102 | 73 | 5.8E-06 | 1.3E-03 |
| | morphology of axons | | | 24 | 16 | 1.3E-05 | 2.1E-03 |
| | potentiation of synapse | -2.913 | 2.475 | 43 | 29 | 3.8E-05 | 5.8E-03 |
| | abnormal morphology of neurites | | | 30 | 23 | 4.2E-05 | 4.1E-04 |
| | development of neurons | -0.955 | 0.531 | 101 | 74 | 4.5E-05 | 2.2E-03 |
| | quantity of neurofilaments | -0.577 | 0.577 | 4 | 4 | 4.6E-05 | 1.6E-05 |
| | axonal transport | -2.425 | | 7 | | 5.2E-05 | |
| | long-term potentiation | -3.148 | 2.767 | 42 | 28 | 6.3E-05 | 9.1E-03 |
| Cortex | quantity of neurofilaments | -0.577 | 0.577 | 4 | 4 | 4.6E-05 | 1.6E-05 |
| | size of axons | | | 4 | 4 | 2.2E-04 | 7.8E-05 |
| | plasticity of neuronal synapse | | | 8 | 7 | 1.6E-04 | 2.2E-04 |
| | plasticity of synapse | -1.664 | 1.026 | 23 | 20 | 4.1E-04 | 2.3E-04 |
| | morphology of dendrites | | | 15 | 16 | 8.8E-03 | 2.4E-04 |
| | cell death of pyramidal neurons | | 1.372 | 4 | 6 | 3.8E-02 | 3.0E-04 |
| | size of neurons | | | 17 | 15 | 6.5E-04 | 3.6E-04 |
| | abnormal morphology of neurites | | | 30 | 23 | 4.2E-05 | 4.1E-04 |
| | morphology of neurites | | | 38 | 30 | 1.3E-04 | 5.0E-04 |
| | long term depression | -1.262 | 0.686 | 17 | 18 | 2.1E-02 | 6.3E-04 |

Activation z-score: 3, 0, -3

**b**

| Overlap with published axonal translatome/transcriptomes | | % overlap | | # of genes | | $log_2$(up/down) |
|---|---|---|---|---|---|---|
| | | Upreg Axons | Downreg Axons | Upreg Axons | Downreg Axons | |
| Translatomes | Adult RGCs, *in vivo* [20] | 2.7 | 3.7 | 30 | 37 | -0.30 |
| | Immature RGCs, *in vivo* [20] | 7.1 | 8.4 | 195 | 220 | -0.17 |
| Transcriptomes | DRG, mature cultures[17] | 10.3 | 4.6 | 271 | 121 | 1.16 |
| | DRG, developing cultures[17] | 13.8 | 6.3 | 331 | 155 | 1.09 |
| | Cortex, mature cultures[19] | 12.9 | 2.4 | 113 | 20 | 2.50 |
| | Cortex, mature cultures, upreg. after injury[19] | 1.3 | 1.2 | 15 | 13 | 0.21 |
| | Cortex, mature cultures, downreg. after injury[19] | 5.5 | 2.9 | 57 | 27 | 1.08 |
| | DRG, injured, developing cultures[18] | 11.6 | 2.4 | 93 | 18 | 2.37 |
| | Motor neurons, developing cultures[46] | 4.1 | 1.1 | 47 | 12 | 1.97 |

% overlap: 15, 7.5, 0

$log_2$(up/down): 1, 0, -1

**Supplementary Figure 7.** a) Functional annotations significantly regulated by learning in the axons and cortex. b) Overlap between genes regulated in axons and published translatomes and transcriptomes in references (16-19).

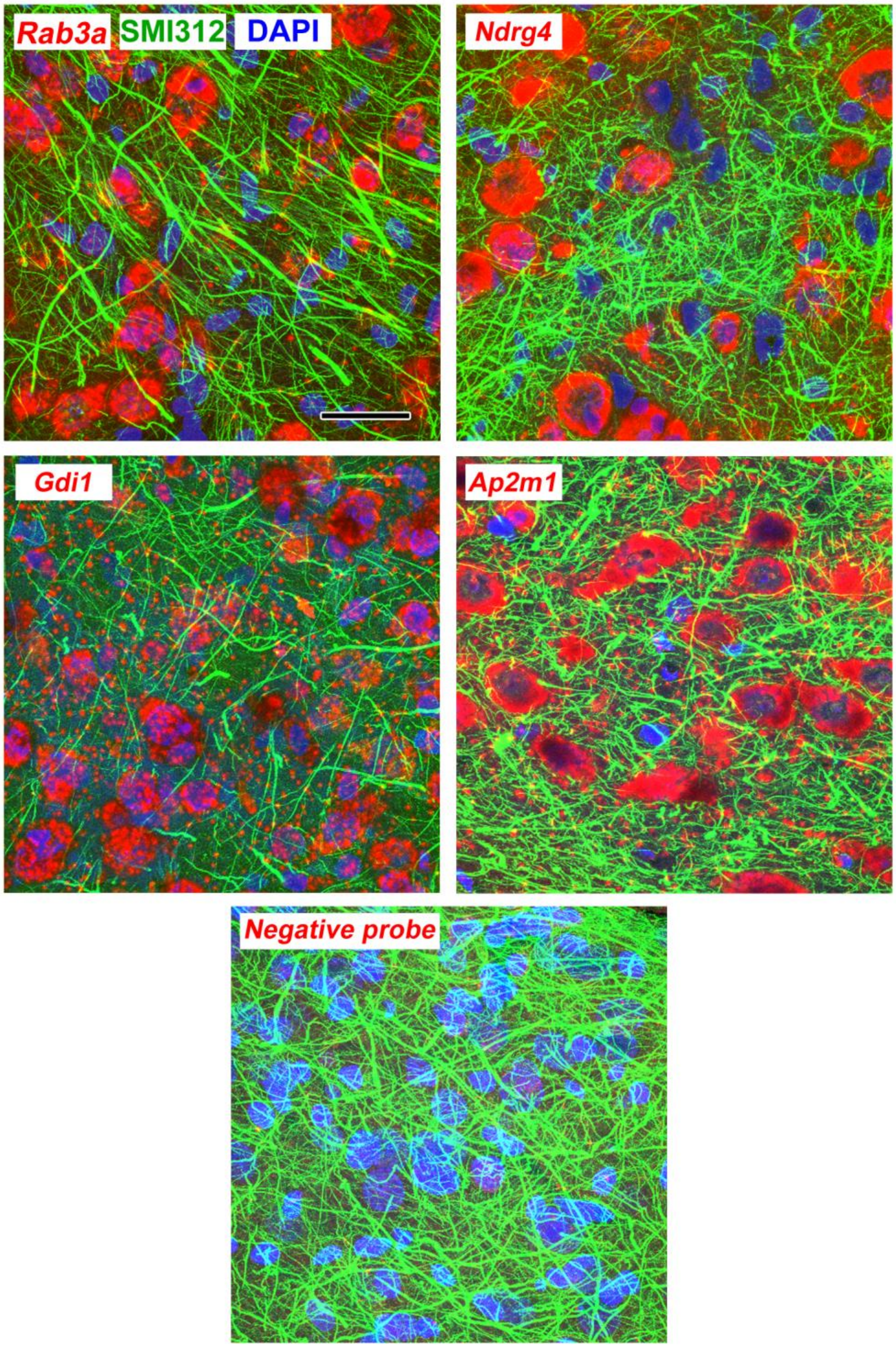


**Supplementary Figure 8. Maximum intensity projections through 3µm (10 confocal images with a 0.3µm z-step size) of lateral amygdala showing FISH labeling and immunolabeling for neurofilaments. Scale = 20 µm.**

**a**

| Gene | transcript | $log_2$(fold change) | adj. p value | contrib. score |
|---|---|---|---|---|
| | | Trained vs control axons | | |
| Snx27 | NM_152847 | -0.99 | 0.021 | 5.99 |
| | NM_001110151 | 0.82 | 0.401 | -4.99 |
| Speg | NM_001108802 | -1.43 | 0.009 | 1.14 |
| | NM_012905 | 0.17 | 0.781 | -0.14 |
| Gng2 | NM_031754 | 0.80 | 0.033 | 2.55 |
| | NM_001257349 | -0.49 | 0.523 | -1.55 |
| | | Control cortex vs axons | | |
| Gria2 | NM_001083811 | 1.53 | 0.001 | 3.78 |
| | NM_017261 | -1.12 | 0.026 | -2.78 |

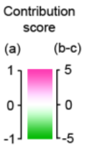


**b**

| Upregulated axons/ downregulated cortex | | contribution score | | | adj. p value | |
|---|---|---|---|---|---|---|
| gene | transcript | axons | cortex | axons-cortex | axons | cortex |
| Apoe | NM_001270681 | 0.00 | 0.01 | -0.01 | 0.876 | 0.942 |
| | NM_001270682 | 0.00 | 0.00 | 0.00 | 0.704 | 0.957 |
| | NM_001270683 | 0.00 | 0.00 | 0.00 | 1.000 | 1.000 |
| | NM_001270684 | 0.75 | 0.73 | 0.02 | 0.031 | 0.730 |
| | NM_138828 | 0.25 | 0.26 | -0.01 | 0.359 | 0.863 |
| Arpp19 | NM_031660 | 0.81 | 0.76 | 0.05 | 0.729 | 0.781 |
| | NM_031660_1 | 0.19 | 0.24 | -0.05 | 0.878 | 0.844 |
| Calm2 | **NM_017326** | 0.32 | 0.27 | 0.05 | 0.002 | 0.018 |
| | **NM_017326_1** | 0.68 | 0.73 | -0.05 | 0.002 | 0.004 |
| Caly | NM_001190399 | 0.37 | 0.13 | 0.24 | 0.588 | 0.983 |
| | NM_138915 | 0.63 | 0.87 | -0.24 | 0.115 | 0.837 |
| Chchd10 | NM_001007008 | 1.00 | 1.00 | 0.00 | 0.004 | 0.327 |
| | NM_001007008_1 | 0.00 | 0.00 | 0.00 | 1.000 | 1.000 |
| Dpy30 | NM_001170545 | 1.00 | 1.00 | 0.00 | 0.005 | 0.550 |
| | NM_173117 | 0.00 | 0.00 | 0.00 | 1.000 | 1.000 |
| Fau | NM_001012739 | 0.59 | 1.04 | -0.46 | 0.432 | 0.499 |
| | NM_001160231 | 0.44 | -0.17 | 0.61 | 0.462 | 0.894 |
| | NM_001160232 | -0.03 | 0.13 | -0.15 | 0.933 | 0.781 |
| Fkbp2 | NM_001134428 | 0.59 | 0.65 | -0.06 | 0.246 | 0.583 |
| | NM_001134429 | 0.41 | 0.35 | 0.06 | 0.286 | 0.680 |
| Gpx4 | NM_001039849 | 0.00 | -0.12 | 0.11 | 0.973 | 0.815 |
| | NM_017165 | 1.00 | 1.12 | -0.11 | 0.335 | 0.716 |
| Hcfc1r1 | NM_001100492 | 1.04 | 1.03 | 0.01 | 0.074 | 0.484 |
| | NM_001185047 | -0.04 | -0.03 | -0.01 | 0.776 | 0.945 |
| Hnrnpa3 | **NM_001111294** | 1.05 | 0.67 | 0.38 | 0.010 | 0.011 |
| | NM_001111295 | -0.02 | 0.00 | -0.02 | 1.000 | 1.000 |
| | NM_198132 | -0.03 | 0.33 | -0.36 | 0.976 | 0.680 |
| Hnrnpd | NM_001082539 | 0.72 | 0.54 | 0.18 | 0.573 | 0.591 |
| | NM_001082540 | 0.71 | 0.27 | 0.44 | 0.409 | 0.720 |
| | NM_001082541 | -0.29 | 0.09 | -0.39 | 0.745 | 0.867 |
| | NM_024404 | -0.13 | 0.10 | -0.23 | 0.907 | 0.943 |
| Klc1 | NM_001081972 | 0.95 | 1.07 | -0.12 | 0.756 | 0.688 |
| | NM_001081973 | -0.09 | -0.03 | -0.07 | 0.930 | 0.908 |
| | NM_001081974 | 0.14 | -0.04 | 0.19 | 0.972 | 0.988 |
| LOC100233176 | NM_001142941 | 0.71 | 2.06 | -1.35 | 0.672 | 0.336 |
| | NM_001142942 | 0.29 | -1.06 | 1.35 | 0.801 | 0.481 |
| Mif | NM_031051 | 1.00 | 1.00 | 0.00 | 0.002 | 0.351 |
| | NM_031051_1 | 0.00 | 0.00 | 0.00 | 1.000 | 1.000 |
| Mrps21 | NM_001126094 | -0.01 | -0.02 | 0.01 | 0.886 | 0.769 |
| | NM_001287116 | 1.01 | 1.02 | -0.01 | 0.156 | 0.680 |
| Naca | NM_001105939 | 0.04 | 0.12 | -0.08 | 0.772 | 0.520 |
| | NM_001198562 | 0.96 | 0.88 | 0.09 | 0.002 | 0.098 |
| | NM_001198580 | 0.00 | 0.00 | 0.00 | 1.000 | 1.000 |
| Ndufv3 | NM_001101011 | 0.87 | 0.82 | 0.05 | 0.185 | 0.697 |
| | NM_022607 | 0.13 | 0.18 | -0.05 | 0.468 | 0.634 |
| Oxr1 | NM_001197332 | 0.82 | 1.13 | -0.31 | 0.086 | 0.009 |
| | NM_001197907 | 0.17 | -0.12 | 0.29 | 0.347 | 0.609 |
| | NM_057153 | 0.01 | -0.01 | 0.03 | 0.903 | 0.941 |
| Park7 | NM_001277249 | 0.00 | 0.00 | 0.00 | 1.000 | 1.000 |
| | NM_001277250 | 0.23 | 0.07 | 0.16 | 0.219 | 0.901 |
| | NM_001277251 | 0.00 | 0.00 | 0.00 | 1.000 | 1.000 |
| | NM_001277252 | 0.74 | 0.83 | -0.10 | 0.014 | 0.337 |
| | NM_001277253 | 0.02 | 0.10 | -0.07 | 0.896 | 0.781 |
| | NM_057143 | 0.01 | -0.01 | 0.01 | 0.691 | 0.649 |
| Pcp4 | NM_001270538 | 0.00 | 0.00 | 0.00 | 0.960 | 0.601 |
| | NM_013002 | 1.00 | 1.00 | 0.00 | 0.004 | 0.337 |
| Pfdn6 | NM_001164718 | -0.01 | 0.67 | -0.68 | 0.996 | 0.738 |
| | NM_212506 | 1.01 | 0.33 | 0.68 | 0.486 | 0.836 |
| Pnkd | NM_001134750 | -0.01 | -0.06 | 0.05 | 0.924 | 0.800 |
| | NM_001134751 | 0.05 | 0.17 | -0.12 | 0.800 | 0.695 |
| | NM_001134753 | 0.96 | 0.89 | 0.07 | 0.357 | 0.591 |
| Rpl35a | NM_021264 | 0.00 | 0.00 | 0.00 | 1.000 | 1.000 |
| | NM_021264_1 | 0.85 | 0.92 | -0.06 | 0.275 | 0.651 |
| | NM_021264_2 | 0.08 | 0.04 | 0.03 | 0.636 | 0.907 |
| | NM_021264_3 | 0.07 | 0.04 | 0.03 | 0.644 | 0.888 |
| | NM_021264_4 | 0.00 | 0.00 | 0.00 | 1.000 | 1.000 |
| Rpl38 | NM_001077592 | 0.00 | 0.00 | 0.00 | 0.761 | 0.843 |
| | NM_001077592_1 | 0.00 | 0.00 | 0.00 | 0.836 | 0.856 |
| | NM_001077592_2 | 1.00 | 1.00 | 0.00 | 0.073 | 0.155 |
| Rps13 | NM_130432 | 0.99 | 0.99 | 0.01 | 0.011 | 0.573 |
| | NM_130432_1 | 0.01 | 0.01 | -0.01 | 0.844 | 0.784 |
| Rps14 | NM_022672 | 0.97 | 0.96 | 0.01 | 0.055 | 0.707 |
| | NM_022672_1 | 0.03 | 0.04 | -0.01 | 0.765 | 0.924 |
| Rps27a | NM_001305443 | 0.33 | 0.27 | 0.06 | 0.188 | 0.602 |
| | NM_031113 | 0.67 | 0.73 | -0.06 | 0.134 | 0.428 |
| Spint2 | NM_001082549 | 0.04 | -0.17 | 0.21 | 0.881 | 0.638 |
| | NM_199087 | 0.96 | 1.17 | -0.21 | 0.216 | 0.645 |
| Tac1 | NM_001124768 | 0.70 | 0.67 | 0.03 | 0.234 | 0.568 |
| | NM_001124769 | -0.02 | -0.05 | 0.03 | 0.956 | 0.967 |
| | NM_001124770 | 0.01 | 0.20 | -0.19 | 0.976 | 0.827 |
| | NM_012666 | 0.31 | 0.18 | 0.13 | 0.515 | 0.835 |
| Tceb1 | NM_001270561 | -0.02 | 0.00 | -0.02 | 1.000 | 1.000 |
| | NM_001270562 | 0.94 | 0.71 | 0.23 | 0.016 | 0.250 |
| | NM_001270563 | 0.05 | 0.11 | -0.07 | 0.849 | 0.669 |
| | NM_022593 | 0.02 | 0.17 | -0.15 | 0.924 | 0.645 |
| Tmem14c | NM_001135169 | 1.10 | 1.20 | -0.10 | 0.144 | 0.487 |
| | NM_134395 | -0.10 | -0.20 | 0.10 | 0.738 | 0.782 |

| Upregulated axons/ downregulated cortex, cont. | | contribution score | | | adj. p value | |
|---|---|---|---|---|---|---|
| gene | transcript | axons | cortex | axons-cortex | axons | cortex |
| Apoe | NM_001270681 | 0.00 | 0.01 | -0.01 | 0.876 | 0.942 |
| | NM_001270682 | 0.00 | 0.00 | 0.00 | 0.704 | 0.957 |
| | NM_001270683 | 0.00 | 0.00 | 0.00 | 1.000 | 1.000 |
| | NM_001270684 | 0.75 | 0.73 | 0.02 | 0.031 | 0.730 |
| | NM_138828 | 0.25 | 0.26 | -0.01 | 0.359 | 0.863 |
| Tpm1 | NM_001034068 | 0.31 | 0.24 | 0.07 | 0.599 | 0.830 |
| | NM_001034069 | 0.01 | -0.03 | 0.03 | 0.842 | 0.704 |
| | NM_001034070 | 0.00 | 0.00 | 0.00 | 1.000 | 1.000 |
| | NM_001034071 | 0.03 | 0.05 | -0.02 | 0.714 | 0.639 |
| | NM_001034072 | -0.08 | 0.13 | -0.21 | 0.875 | 0.880 |
| | NM_001034073 | 0.19 | 0.26 | -0.07 | 0.561 | 0.678 |
| | NM_001034074 | -0.06 | 0.08 | -0.14 | 0.671 | 0.510 |
| | NM_001034075 | 0.20 | -0.06 | 0.26 | 0.548 | 0.937 |
| | NM_001301336 | -0.04 | -0.14 | 0.09 | 0.774 | 0.680 |
| | NM_001301342 | -0.03 | -0.01 | -0.02 | 0.860 | 0.956 |
| | NM_001301736 | 0.46 | 0.06 | 0.40 | 0.715 | 0.990 |
| | NM_019131 | 0.01 | 0.40 | -0.39 | 0.988 | 0.928 |
| Uqcr11 | NM_001126097 | 0.95 | 0.98 | -0.03 | 0.004 | 0.241 |
| | NM_001287109 | 0.05 | 0.02 | 0.03 | 0.779 | 0.951 |

**c**

| Downregulated axons/ upregulated cortex | | contribution score | | | adj. p value | |
|---|---|---|---|---|---|---|
| gene | transcript | axons | cortex | axons-cortex | axons | cortex |
| Akap2 | NM_001011974 | 0.21 | 0.28 | -0.08 | 0.135 | 0.724 |
| | NM_001309260 | 0.79 | 0.72 | 0.08 | 0.002 | 0.137 |
| Begain | NM_001111115 | 0.07 | 0.00 | 0.07 | 0.833 | 1.000 |
| | NM_024163 | 0.93 | 1.00 | -0.07 | 0.121 | 0.004 |
| Camk2b | NM_001042354 | -0.01 | 0.01 | -0.02 | 1.000 | 0.596 |
| | NM_001042356 | 0.19 | -0.17 | 0.37 | 0.809 | 0.834 |
| | NM_021739 | 0.81 | 1.16 | -0.35 | 0.428 | 0.057 |
| Csnk1g2 | **NM_001033870** | 0.83 | 0.77 | 0.06 | 0.022 | 0.004 |
| | NM_023102 | 0.17 | 0.23 | -0.06 | 0.503 | 0.283 |
| Dlgap1 | **NM_001304287** | 1.01 | 0.91 | 0.11 | 0.005 | 0.009 |
| | NM_022946 | -0.01 | 0.09 | -0.11 | 0.931 | 0.728 |
| Dlgap3 | NM_001276304 | 0.07 | 0.12 | -0.04 | 0.753 | 0.724 |
| | NM_001301876 | 0.93 | 0.88 | 0.04 | 0.028 | 0.199 |
| Iqsec2 | NM_001277386 | -0.01 | -0.04 | 0.03 | 1.000 | 1.000 |
| | NM_001277425 | 1.01 | 1.04 | -0.03 | 0.054 | 0.129 |
| Kdm5a | **NM_001277177** | 1.00 | 1.00 | 0.00 | 0.006 | 0.006 |
| | NM_001277178 | 0.00 | 0.00 | 0.00 | 1.000 | 1.000 |
| Mbp | NM_001025289 | 0.00 | 0.00 | 0.00 | 0.909 | 0.976 |
| | NM_001025291 | 0.04 | 0.02 | 0.02 | 0.547 | 0.747 |
| | NM_001025292 | 0.15 | 0.20 | -0.05 | 0.161 | 0.033 |
| | NM_001025293 | 0.13 | 0.11 | 0.02 | 0.070 | 0.549 |
| | NM_001025294 | 0.02 | 0.02 | 0.00 | 0.817 | 0.798 |
| | **NM_017026** | 0.66 | 0.65 | 0.02 | 0.002 | 0.004 |
| Mprip | NM_001034022 | 0.95 | 0.92 | 0.02 | 0.002 | 0.004 |
| | NM_053814 | 0.05 | 0.08 | -0.02 | 0.186 | 0.114 |
| Nexn | NM_139230 | 0.86 | 0.94 | -0.09 | 0.002 | 0.231 |
| | NM_139231 | 0.14 | 0.06 | 0.09 | 0.468 | 0.936 |
| Nsmf | NM_001270626 | 0.57 | 0.00 | 0.57 | 0.543 | 1.000 |
| | NM_001270627 | -0.17 | 0.34 | -0.51 | 0.757 | 0.758 |
| | NM_001270628 | -0.10 | 0.08 | -0.18 | 0.644 | 0.744 |
| | NM_057190 | 0.46 | 0.04 | 0.43 | 0.330 | 0.803 |
| | NR_073057 | 0.24 | 0.54 | -0.30 | 0.699 | 0.765 |
| Pde2a | NM_001143847 | 0.10 | -0.01 | 0.11 | 0.633 | 0.990 |
| | NM_001270604 | 0.65 | 0.63 | 0.02 | 0.481 | 0.679 |
| | NM_031079 | 0.25 | 0.38 | -0.13 | 0.821 | 0.805 |
| Pip5k1c | NM_001009967 | 0.61 | 0.53 | 0.08 | 0.600 | 0.184 |
| | NM_001033970 | 0.39 | 0.47 | -0.08 | 0.715 | 0.373 |
| Plec | **NM_001164296** | 0.84 | 0.85 | -0.01 | 0.002 | 0.004 |
| | NM_001164297 | 0.01 | 0.12 | -0.10 | 1.000 | 0.708 |
| | NM_001164298 | 0.04 | 0.01 | 0.02 | 1.000 | 1.000 |
| | NM_001164299 | 0.00 | 0.00 | 0.00 | 1.000 | 1.000 |
| | NM_001164302 | 0.00 | 0.00 | 0.00 | 1.000 | 1.000 |
| | NM_001164303 | 0.00 | 0.00 | 0.00 | 1.000 | 1.000 |
| | NM_001164304 | 0.07 | 0.00 | 0.07 | 0.739 | 1.000 |
| | NM_001164305 | 0.04 | 0.00 | 0.04 | 0.710 | 1.000 |
| | NM_001164307 | 0.00 | 0.00 | 0.00 | 1.000 | 1.000 |
| | NM_001164308 | 0.00 | 0.02 | -0.02 | 1.000 | 1.000 |
| | NM_022401 | 0.00 | 0.00 | 0.00 | 1.000 | 1.000 |
| Prkcb | NM_001172305 | 0.21 | 0.37 | -0.15 | 0.726 | 0.520 |
| | NM_012713 | 0.79 | 0.63 | 0.15 | 0.050 | 0.091 |
| Rasgrf1 | NM_001105753 | 0.00 | -0.01 | 0.01 | 1.000 | 0.822 |
| | NM_001170531 | 1.00 | 1.01 | -0.01 | 0.005 | 0.070 |
| Ryr2 | **NM_001191043** | 0.97 | 0.94 | 0.03 | 0.005 | 0.004 |
| | NM_032078 | 0.03 | 0.06 | -0.03 | 1.000 | 0.720 |
| Shank2 | NM_001004133 | 0.47 | 0.84 | -0.37 | 0.140 | 0.487 |
| | NM_133440 | 0.01 | 0.04 | -0.04 | 1.000 | 1.000 |
| | NM_133441 | 0.14 | 0.23 | -0.08 | 0.515 | 0.565 |
| | NM_201350 | 0.38 | -0.11 | 0.49 | 0.171 | 0.909 |
| Slc25a3 | NM_001270788 | 0.00 | 0.00 | 0.00 | 0.944 | 1.000 |
| | **NM_139100** | 1.00 | 1.00 | 0.00 | 0.018 | 0.004 |

**Supplementary Figure 9.** Transcript-level analysis. The contribution score (change in FPKM transcript/change in FPKM gene) indicates the effect of learning on a transcript relative to the net effect on all transcripts of the same gene, with a negative score indicating differences in opposite directions between the transcript and gene. Adjusted p-values for each transcript are highlighted at <0.05. a) Three transcripts were found to be regulated by learning in the axons that were not differentially expressed at the gene level. In each case, a second transcript was affected non-significantly in the opposite direction. The two transcripts of *Gria2* were differently distributed in the control group, with one enriched in axons and the other in cortex. b-c) Genes regulated in both axons and cortex (b; upregulated in axons/downregulated in cortex, c; downregulated in axons/upregulated in cortex) with multiple transcripts in the dataset. The difference between the score in the axons and cortex ("axons – cortex") indicates the degree of asymmetry, with positive numbers indicating transcripts which were affected proportionally more in the axons than cortex. Values near zero indicate transcripts that were similarly affected in both areas. Transcripts with significant effects in both areas are shown in bold type.
